# Ordering from the edge: Chiral multicellular pattern formation directed by actin self-organisation in cells

**DOI:** 10.1101/2025.10.14.682394

**Authors:** Wenzheng Shi, Dinh Thach Lam Nguyen, Wei Jia Goh, Hui Ting Ong, Run Bin Tan, Chaoyu Fu, Alexander D. Bershadsky, Alex Mogilner, Yee Han Tee

**Affiliations:** Courant Institute and Biology Department, New York University, New York, NY, USA; Mechanobiology Institute, National University of Singapore, Singapore, Singapore; Department of Physics, National University of Singapore, Singapore 117542, Singapore; Department of Molecular Cell Biology, Weizmann Institute of Science, Rehovot 7610001, Israel

## Abstract

The mechanisms underlying both the establishment of mirror symmetry and deviations from it in development of bilateral multicellular organisms remain insufficiently understood. Actin cytoskeletons of individual cells exhibit intrinsic chirality, and a strong correlation exists between single-cell actin fibres chiral organisation and the collective alignment of cells confined to rectangular adhesive islands (2D-microtissues). Here, we demonstrate how multicellular chiral patterns can be inferred from the chiral behaviour of actin fibres in individual cells. By analysing chiral actin systems in cells with elliptical and semi-circular shapes, representing inner and boundary positions within 2D-microtissues, we defined the rules of chiral motile behaviour and formulated two models of cell alignment: (i) chiral rotation of inner cells and (ii) chiral tilting of boundary cells relative to island edges. In both models, neighbouring cells are also mutually aligned. Systematic variation of island area and aspect ratio, combined with dynamic observations, revealed the primary role of boundary cells. Chiral order first emerged at tissue boundaries and then propagated inward. This outside-in mechanism also explains why mirror-symmetric cell groups are difficult to achieve with uniformly chiral cells, but can be obtained either by reversing chirality in one subgroup or by enlarging the microtissue to minimise boundary influence.

## Introduction

Asymmetric biological structures, from molecules to organisms, underlie life as we know it. The asymmetry type that has received most of attention is polarity, the difference between ‘head’ and ‘tail’. Many mechanisms of relevant symmetry breaking were discovered, from stereospecific linear assembly of cytoskeletal polymers to Turing patterns in reaction-diffusion systems. In bilateral organisms characterised by mirror symmetry between two halves of the body, there is another biological asymmetry – deviations from the mirror symmetry known as chirality^1-8^. While in some organisms like snails^9, 10^ or fiddler crab^11^ chirality is manifested in the prominent asymmetry of their body shapes, majority of multicellular bilateral organisms preserve external mirror symmetry of the body but positioning and organisation of some inner organs like heart, brain and gut are chiral. In spite of obvious biological importance, molecular and cellular mechanisms of both organismal mirror symmetry and deviation from it are insufficiently understood.

Many chemical compounds can exist as chiral isomers (enantiomers) and it is known since 19^th^ century that all organisms demonstrate total “enantio-selectivity” being built of L-amino acids, D-sugars, etc even though some interesting exception are possible^12^. It is not known however if enantio-selectivity is related to processes of chiral morphogenesis outlined above. The breakthrough in understanding of organismal chirality emerged from realising that multiple cell types demonstrate built-in chirality based on asymmetric self-organisation of the cytoskeleton.

A very important and well-studied pathway for establishing of organismal chirality is operating through activity of cilia in cells of a special transient “left-right organiser” known as node (in mouse) or Kupffer’s vesicle (in zebrafish). These cilia create an asymmetric flow which is detected by the immotile sensory cilia of surrounding cells which in turn initiates a signalling cascade resulting in establishing left-right asymmetry^13-16^. The cilia are cytoskeletal organelles rooted in the bulk of the cell cytoskeleton and it is not clear whether the initial asymmetry of the flow is developed due to asymmetric organisation of cilia itself or due to its association with other cytoskeletal asymmetric structures. Most striking is a recent observation that in species like birds and reptiles the cells in the node homologue, known as Hensen’s node, lacked the cilia and the asymmetry is developed by coordinated movements of cells comprising an “active torque dipole” ^17-19^. These cellular activities most probably depend on the actomyosin cytoskeleton.

Indeed, several seminal observations of chirality development in cell collectives *in vitro*^20, 21^ suggested that development of the chiral pattern depends on the actin cytoskeleton function. In our previous studies, we described the chiral self-organisation of the actin cytoskeleton in individual fibroblast-type cell under isotropic confinement^22^ and identified several actin-associated proteins regulating actin filament polymerisation and crosslinking involved in this process^23^. In invertebrate development, some actin-related proteins were shown to be involved in establishment of left-right asymmetry of *Drosophila* gut and genitalia ^24-27^, the chiral morphology of pond snails^28-30^ and the asymmetric rotation of *Caenorhabditis elegans* zygote^31^. Moreover, the heart looping in vertebrate embryos seems to involve nodal-independent actin-driven processes^32, 33^.

In general, the origin of actin-dependent chirality is the helical chiral organisation of individual actin filaments. The mechanism converting the chirality of individual filament into chirality of higher order actin structures and eventually the chirality of entire cell remains an active area of research. Among proposed mechanisms are the chiral rotation of the actin filament during its polymerisation by formins or chiral torque developed via interaction between actin filaments and different myosins or a combination of these^22, 31, 34-39^. Even though the exact functioning of such mechanisms is not completely clear, the phenomenology of the chiral self-assembly of the actin cytoskeleton in individual circular cells is described in considerable detail and effects of various experimental interventions are explored^23^. Furthermore, it appears that there is a strong correlation between ability of individual circular cells to form an actin cytoskeleton in a chiral fashion and the establishing of chiral pattern in the groups of such cells confined to rectangular adhesive islands^23^. This correlation obviously deserved explanation and analysis as a possible clue to understanding of the basic mechanisms of morphogenesis in the simple tissue-like systems (2D-microtissues).

Computational modelling is an invaluable tool that can accelerate understanding by being able to rapidly examine relative importance and interdependence of molecular mechanisms. Modelling was used earlier to explain the emergence of collective cellular chiral alignment from the chiral behaviour of individual cells. Hydrodynamic-like theories, as well as discrete mechanical models treating cells as elongated particles showed that chiral interactions of cells with boundaries and each other can lead to chiral collective patterns^40-43^. Comprehensive review of mathematical models of individual and collective cell chirality can be found in Rahman *et al*.^44^. Of note, the chirality of individual cells in majority of these studies was usually introduced just as a parameter characterising nematic behaviour of cells rather than derived from the chiral self-organisation of the actin cytoskeleton.

In the present study, we first carefully examined the actin self-organisation in the cells with the shape resembling that of cells in the confined monolayer. In particular, we focused on two types of cell morphology: the semi-circular cells mimicking the cells starting to align along the boundaries of the adhesive islands and the elliptical cells representing the bulk of inner cells of the islands. We found that the actin cytoskeleton self-organised in a chiral manner in both these morphological types and determine the mode of cell shape change. As a result, the elliptical (quasi-inner) cells demonstrated chiral rotation, while the semi-circular (quasi-boundary) cells developed a chiral tilt relatively to the boundary. Based on these experimental findings, we formulated the rules describing the chiral behaviour of the inner and boundary cells in the 2D-microtissues. These rules, together with a natural assumption about the mode of cell alignment permitted us to propose two basic models of multicellular pattern development. In the first model, the chiral pattern formation emerged from rotation and alignment of inner cells that propagate from the inner area to the boundary of the microtissue. The second model suggests that chiral alignment originate from the tilting of boundary-interacting cells that propagate from the boundary inward.

Surprisingly both models satisfactorily described the development of chiral pattern of cell alignment in our standard experimental model, human fibroblast cells confined to 300 × 600 μm^2^ rectangular adhesive island. However, the varying of micropattern area and aspect ratio revealed that the predictions of boundary-cell-driven model demonstrated significantly better agreement with the experimental data than those of the inner-cell-driven model. We then confirmed the boundary-cell chirality model predictions by quantifying dynamic evolution of the collective chiral behaviour, observing that indeed the chiral order emerged first at the periphery of the rectangular microtissue and only later established in its inner area. Thus, these simple experimental models revealed a potentially important principle of multicellular organisation in the tissues attracting the attention to the cellular morphogenetic activities at their boundaries.

Finally, based on this theory, we started to analyse a newly emerged basic question: how the mirror-symmetrical tissues and organs can be built from the cells with intrinsic chirality. In a simple toy model we considered the alignment of intrinsically chiral fibroblasts inside mirror-symmetrical parallelogram-shaped patterns. The surprising conclusion was that due to the clash between intrinsic cell chirality and chirality of the parallelogram-shaped boundary, the mirror-symmetrical parallelograms filled with the cells lost their mirror symmetry. This prediction, fully confirmed by our experiments, emphasised the non-triviality of the process of formation of mirror symmetrical cellular structures in development. We then outlined the possible mechanisms that can secure formation of mirror symmetrical complexes from the chiral cells.

## Results

### Chiral behaviour of stress fibres

Previously, we found a strong correlation between the chirality of the actin cytoskeleton in individual cells confined to circular adhesive pattern and the emergence of chiral pattern in cell groups^23^ (see Fig. 1a). Within a cell collective, cells adopted an elongated (spindle-shape or semi-circular) rather than a circular morphology. Chiral pattern of cell groups confined to rectangular islands is manifested by preferential orientation of inner cells along the bottom-left-to-top-right diagonal, making inverted letter N (looking like Cyrillic И)^23^. At the same time, the cells spreading in the proximity of the boundary of the rectangular island usually acquired an approximately semi-circular shape (Fig. 1b). To understand how the chiral organisation of actin in a single cell can drive chiral alignment within a cell group, we investigated how the system of actin fibres self-organise in cells confined to an elliptical shape (representing inner cells), as well as to semi-circular shape (representing cells spreading at the boundary between adhesive and non-adhesive areas). We also examined how the actin fibres organisation changes upon release of the cells from these patterns.

**Figure 1.**
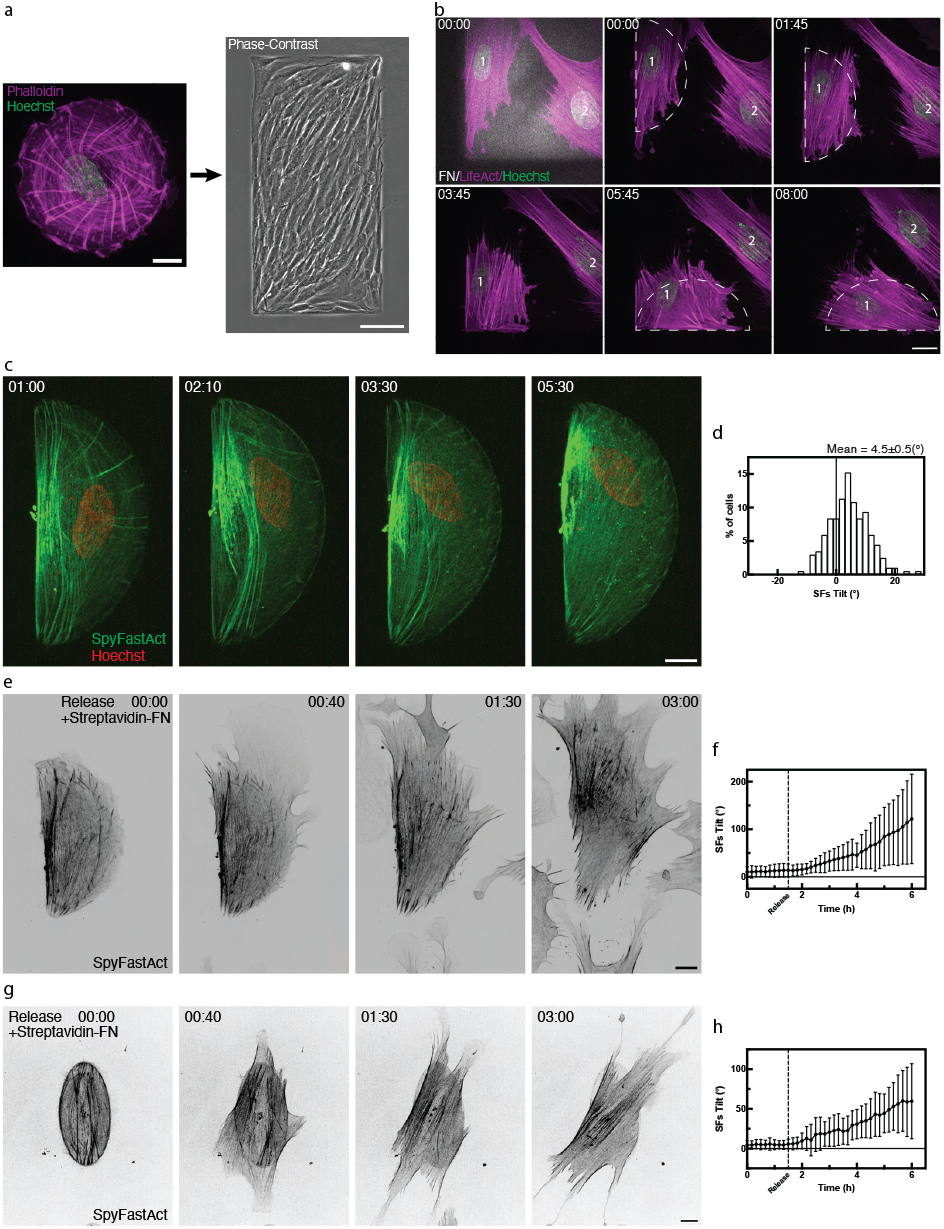
Chiral behaviour of stress fibres in individual confined cells. **a**, (Left) Typical example of chiral organisation of the actin cytoskeleton (magenta) in a single cell plated on a circular fibronectin island (48 μm diameter). Actin was labelled by phalloidin, and the nucleus was labelled by Hoechst-33342 (green). (Right) Phase-contrast image of multicellular chiral alignment pattern, resembling the Cyrillic letter “И”, on a rectangular fibronectin island (300×600 µm) after 48 h. Scale bars: 10 μm (left), 100 μm (right). **b**, Time-lapse imaging of a LifeAct-transfected cell (magenta) interacting with the boundary of a fibronectin-coated (FN; white) rectangular island (cell 1, dashed semi-circular outline). Only selected regions of the rectangular island could be visualised by high magnification live-cell imaging; timepoints (hh:mm) started within 1 h post-cell seeding. Nuclei 1 and 2 (green) were labelled by Hoechst-33342. Note that the stress fibres in these cells are tilted relative to the boundary. Scale bar: 20 μm. **c**, Time-lapse sequence showing the dynamic self-organisation of the actin cytoskeleton (green) and associated nuclear (red) movement in a cell plated on a semi-circular fibronectin island (radius 40 μm; area 2500 μm^2^) at selected time points (hh:mm). Actin and nucleus were visualised using SpyFastAct-650 and Hoechst-33342 respectively. Scale bar: 10 μm. See also Extended Data Fig. 1b, Extended Data Fig. 2a,b and Supplementary Video 1. **d**, Histogram showing the distribution of stress fibres (SFs) tilt angles in fixed cells 6 h after plating on semi-circular fibronectin island and labelled with phalloidin. Mean ± s.e.m. value for n = 204 cells is indicated at top right. See also Extended Data Fig. 1a. **e, g**, Time-lapse sequence showing actin dynamics (visualised by SpyFastAct-650) and deformations of cells after release from semi-circular (**e**) or elliptical (**g**) confinement. Timepoints (hh:mm) begin after release was induced by the addition of streptavidin-fibronectin to PLL-PEG-biotin passivated dishes. Scale bars: 10 μm. See also Supplementary Videos 2,3. **f, h**, Quantification of the mean stress fibre tilt angle over time for cells released from semi-circular (**f**, n = 26 cells) and elliptical (**h**, n = 28 cells) micropatterns. Confinement was released at t = 1.5 h. Data are presented as mean ± s.d.

Actin radial fibres in confined cells grow from the focal adhesions located at the cell boundary^22^ (Extended Data Fig. 1). On circular islands, these adhesions are evenly distributed along the edge of the circles, while on both elliptical and semi-circular islands, their distribution strongly depend on the curvature of the boundary (in line with previous studies^45-47^). In particular, on elliptical islands, majority of focal adhesions concentrated in proximity of the ellipse vertices^23^, while on semi-circular islands, focal adhesions are larger and more numerous at the endpoints of the diameter (in the ‘corners’), but also form along the arc, yet significantly less developed along the straight side (diameter) (Extended Data Fig. 1).

Thus the radial fibres in cells on elliptical substrate were more pronounced at the vertex region^23^ (Fig. 1g), while on semi-circular substrate they were radiating from the endpoints of the diameter (Fig. 1c,e and Extended Data Fig. 1a). Similarly to the radial fibres in cells on circular substrate^22^, the radial fibres growing from the focal adhesions on elliptical and semi-circular micropatterns turned clockwise during their growth (as viewed from the medium towards the substrate) (Fig. 1c). In elliptical cell such tilting culminates by the diagonal orientation of the stress fibres^23^ (Extended Data Fig. 1c). In semi-circular cell, the outcomes of clockwise tilting at either ends of the diameter differ. Those radial fibres which tilt away from the diameter eventually occupy the entire cell area and together with the fibres emerging from the arc of the semi-circle formed the final actin array (Fig. 1c and Extended Data Fig. 1a,b and Supplementary Video 1). At the same time, the fibres emerging from the opposite end of the diameter, tilted towards the diameter and contributed to the formation of actin bundles oriented along the diameter. As a result, the array of stress fibres in semi-circular cell will radiate from a single end of the diameter (Fig. 1c and Extended Data Fig. 1a,b). The final mean orientation of the stress fibres is several degrees (°) to the right relative to the semi-circle diameter (Fig. 1d and Extended Data Fig. 2a,b), similar to the chiral deviation of stress fibres from the ellipse’s long axis as previously reported^23^.

These clockwise-tilted stress fibres orientation in both the semi-circular and elliptical micropatterns is preserved upon releasing cells from confinement by coating the surrounding non-adhesive area with fibronectin as described in Isomursu *et al*.^48^(Supplementary Videos 2,3). Moreover, following the release, cells elongated in the direction of their stress fibres, forming new lamellipodia in the proximity of the fibres ends (focal adhesions) (Fig. 1e,g and Extended Data Fig. 2c,f,e,h). The rotation of the stress fibres continued in the same direction inside the new cell shape, further increasing the average tilt of the stress fibres (Fig. 1f,h and Extended Data Fig. 2d,g). Then, the new lamellipodia formed following the same rule, resulting in apparent rotation of the cell long axis. Obviously, the configuration of cell outlines upon release from confinement becomes more erratic with time (Supplementary Videos 2,3).

### “Boundary-cell” and “Inner-cell” Models

Thus, asymmetric rotation of the stress fibres, similar to that observed previously in cells confined to circular pattern, plus the rule of formation of new lamellipodia in proximity of the ends of stress fibres observed experimentally, permit to explain the asymmetric deformations of individual cells: (i) chiral rotation of elliptical cells and (ii) tilting of cells originally aligned along the boundary of the adhesive island relatively to this boundary. To explain the behaviour of cell collective, we can propose two main hypotheses based on the observations described above (Fig. 2).

**Figure 2.**
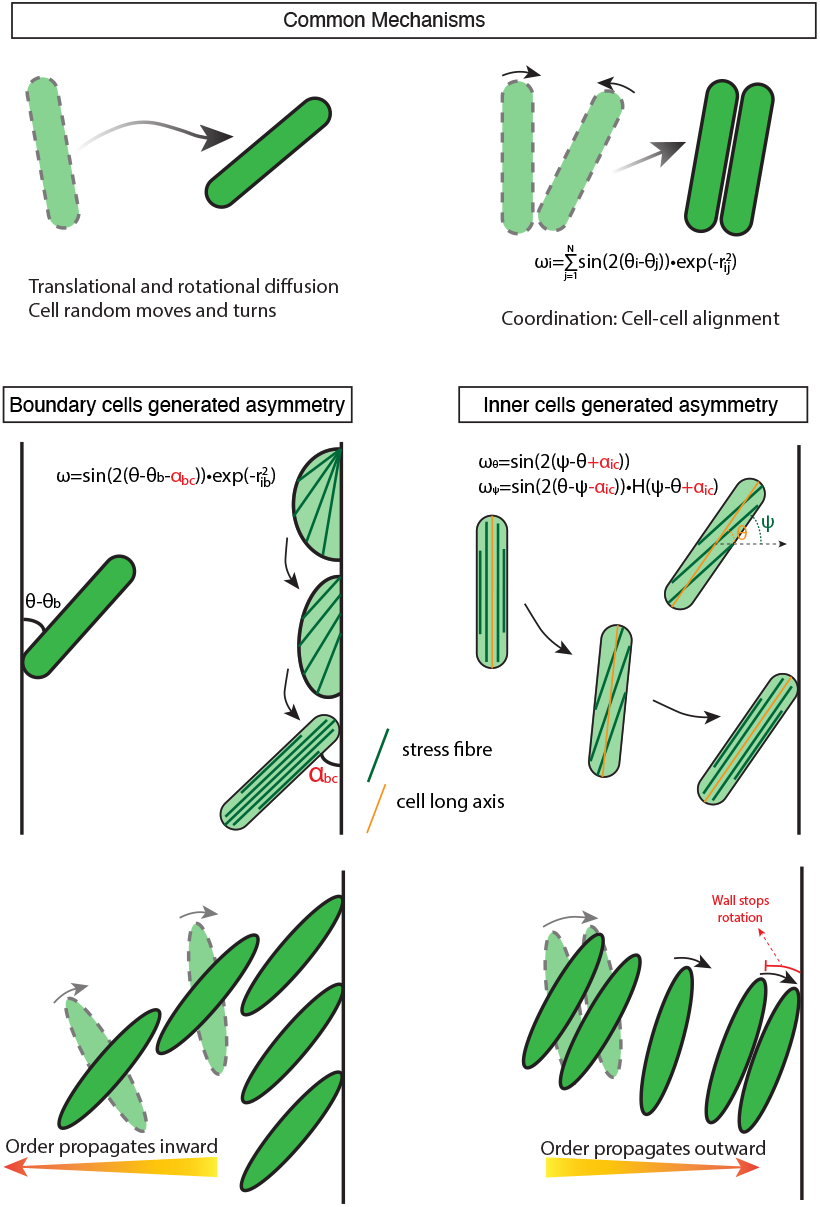
“Boundary-cell” and “Inner-cell” chirality models. The top of the figure demonstrates the common mechanisms shared by both models: (Left) translational and rotational diffusion of individual cell; (Right) alignment interaction between cell pairs, inducing angular velocity differences that tend to synchronise neighbouring cells by coupling their orientation differences. The angular velocity imposed on cell j by all other cells are shown in the figure. Here N is the total number of cells, r_ij_ is the distance between cell i and cell j. The alignment between cell and boundary following the same dynamic as cell-cell interactions. r_ib_ is the distance between cell i and the wall and *θ*_*b*_ is the angle parallel to the wall. The middle of the figure shows the assumptions specific to two models. At the left is the boundary-cell model where cell-boundary interaction is modified by imposing the tilt angle *α*_bc_. At the right is the inner-cell model where orientations of the cell long axis θ and stress fibres ψ are two coupled dynamic variables. *α*_ic_ is the persistent offset angle between stress fibres and the cell axis. H is the Heaviside function, enforcing clockwise rotation of the stress fibres. The bottom panel represents the direction of chirality propagation. In the boundary-cell model, the chirality emerges at the boundary and gradually propagates inward through cell-cell alignments. In the inner-cell model, the chirality arises inside the island from individual cell rotations and mutual alignments. Consequently, the cells rotate collectively until the boundary-cell interaction stalls the rotation resulting in the collective chiral tilt.

Both treat individual cells as elongated objects that we depict as ellipsoids, but in the simplified models not the actual complex shape characterises the cell but two variables – the position of its centre and orientation of its long axis. In both models, in agreement with the experimental observations, (1) the cells move, and their long axis turn randomly on the 2D adhesive island; (2) neighbouring cell pairs tend to align with each other; and (3) cell near the boundary tend to align with the boundary. We simulated the model with just these three rules, and found, expectedly, that if the alignment strength is great enough compared to the magnitudes of random movements and turns, then the cell groups tend to align with each other and with one of the diagonals of the rectangular island with equal frequency – the latter, based on simulations, is explained by the emergent tendency of the whole group to maximise its collective long axis (Extended Data Fig. 3). Hence, additional assumptions beyond these three rules are required to explain the asymmetric cell alignment within rectangular confinements.

In the first, boundary-cell model, there is just one additional phenomenon: the cells that are near the boundary tend to tilt by a certain small angle relative to the boundary, as observed. In this model, this intrinsic chirality bias in the cell-boundary interactions propagates inward generating the collective chiral pattern (Fig. 2).

In the second, inner-cell model, each cell undergoes an independent, slow clockwise rotation, reflecting the observed process: in the elongated cells, the stress fibres slowly turn clockwise relative to the cell axis, then the cell shape slowly adjusts its axis to the stress fibres direction, and so on (Fig. 2). Effectively, the edge cells become tilted, with the tilt determined by the mechanical balance between the individual rotation and the cell-boundary alignment. In this model, the collective chirality in a way propagates from inside the island where the cells organise into aligned, slowly rotating, groups to the boundaries. The models’ equations are explained in Methods. There are very few model parameters. The cell numbers, tilt angles and random movement/turning rates are informed by the experiments. The alignment magnitudes are found from fitting the model predictions to the experimental data described in the next section. After that, there is no freedom left – all parameters are fixed, and several other experiments truly test the models.

### Diversity of orientation patterns stems from a ‘competition’ of the long and short sides of the island and can be explained by both models

Cells confined to rectangular adhesive islands (2D-microtissues) displayed several characteristic types of patterns. To classify them, each 2D-microtissue can be characterised by an array of 8×16 nematic directors (red in Fig. 3a), with each director reporting the average local orientation of the cells (Fig. 3a). To qualitatively categorise different orientation patterns, we divided individual microtissue into four zones, each comprising an 8×4 nematic director array, and calculated the mean orientation for each zone, which was used to determine its orientation pattern (Fig. 3a and Extended Data Fig. 4a). Thus, each zone was qualitatively classified as right-tilted (/), left-tilted (\), vertical (|) or horizontal (–) (Extended Data Fig. 4b) and entire microtissue was categorised by an ordered combination of four such symbols (Fig. 3a). We succeeded to classify all observed patterns into six classes denoted as “И”, “N”, “C”, “Z”, “S” or “I” as shown in Extended Data Fig 4c. Among the observed patterns, the “И”-orientation was predominant (∼51%), while “N”, “C”, and “I” orientations each appeared at approximately 14%, 30% and 1%, respectively. The “Z”-orientation occurred less than 2%. Notably, “S” the mirror-image counterpart of chiral “Z”-orientation was almost absent in experimental control populations.

**Figure 3.**
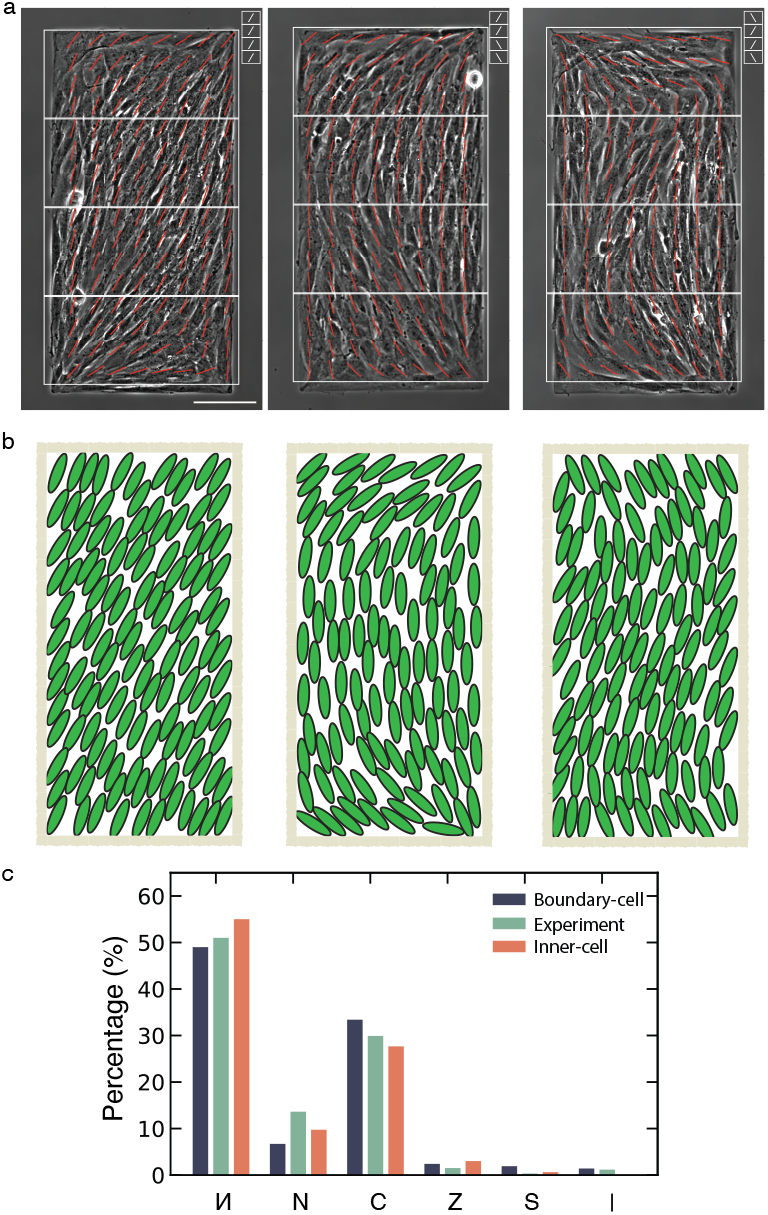
Diverse cell orientation patterns can be reproduced by simulation. **a**, Phase-contrast images of fibroblast 2D-microtissues on rectangular micropatterns 48 h after seeding. The average local cell orientation (nematic director) is indicated by red lines. To classify the overall orientation pattern, each microtissue was divided into four quadrants (white boxes), and the dominant tilt direction in each was assessed (insets: /, right-tilted; \, left-tilted). Representative “И” (left), “C” (middle), and “Z” (right) patterns are shown. Scale bar, 100 µm. See Methods for classification details and Extended Data Fig. 4. **b**, Representative simulation outputs showing the emergence of the orientation patterns observed in **a**. Individual cells are depicted as green ellipses and the micropattern boundary is outlined in light brown. **c**, Frequency distribution of the six orientation patterns observed in experiments (green bar) and predicted by two distinct simulation models – boundary-cell (dark blue bar) and inner-cell (orange bar). n= 602 microtissues for experiments; n = 2,000 for each simulation condition.

We successfully fitted the boundary-cell model to experimentally determined percentages of each class of pattern by varying a parameter characterising the relative strength of cell-to-cell and cell-to-boundary alignment (see Fig. 3b and Methods). According to the model, this observed diversity of the collective orientational patterns stems from the peculiar ‘competition’ of the long (‘vertical’) and short (‘horizontal’) sides of the island for the cells: near the long side, cells tend to orient ‘bottom left’ to ‘top right’, but near the short side, cells tend to orient ‘bottom right’ to ‘top left’. The most frequent “И”-pattern evolves when two long sides ‘win’ over two short sides. The next dominant “C”-pattern emerges when one long and one short side win; rarer “N”-pattern – when two short sides win. Clearly, the frequency of a certain pattern correlates with the length of the fraction of the boundary informing such pattern. “Z”-pattern appears when all four boundaries of the rectangle win, but locally, not globally. The “S”-pattern seemingly frustrates all four boundary conditions, and so is observed very rarely. Also very rare “I”-pattern ‘ignores’ all boundary conditions. According to the model, any of these patterns are quasi-steady states that result from competition between tendencies of all cells to align with each other and frustrations of conflicting alignments with different island boundaries and corners and emerge depending on random initial conditions and random turns and shifts of individual cells. Expectedly, similar fitting of the inner-cell model also demonstrated a good agreement with experimental data.

### Orientation patterns in 2D-microtissues confined to rectangular adhesive islands of varying area rule out the inner-cell model and support the boundary-cell model

To further compare experimental results with predictions of both models, we varied the areas of rectangular adhesive islands (without changing their aspect ratios) and calculated the distributions of average island nematic director values for rectangles with different areas (Fig. 4a). Then we computed these distributions for boundary-cell and inner-cell models (Fig. 4b,c). It is clearly seen from comparison of histograms in Fig. 4a to those in Fig. 4b and 4c that the boundary-cell model better fits the experimental distribution, whereas the inner-cell model does not. The qualitative explanation is: in the boundary-cell model, the tilt enforced by the boundary cells propagates only to a finite distance into the bulk of the cell group because of the random turnings and movements. So, on the larger islands, the cells near the centre do not ‘feel’ the boundary effects. On the other hand, in the inner-cell model, all cells in the centre align first, and then their collective rotation is stopped by the boundaries – in this mechanism, the greater the islands, the larger the tilted angle. To quantify these observations and simulations, we introduced a parameter called chiral index as the ratio between fractions of right-tilted to left-tilted nematic directors i.e. the ratio between dark green and yellow areas in histograms in Figs. 4a-c. Plotting the experimentally determined chiral indices against the outer-to-inner area ratio of different rectangles (defined as the ratio between the 50 μm-wide peripheral zone and the inner region) reveals a marked discrepancy with the predictions of the inner-cell model, but a strong agreement with those of the boundary-cell model (Fig. 4d). To support this result by further experiments, we also varied the aspect ratio of the rectangles and demonstrated that the boundary-cell model is in a very good agreement with this set of experimental data (Extended Data Fig. 5). Note that there is neither experimentally observed (Extended Data Fig. 5a), nor model-predicted chirality on the square islands (Extended Data Fig. 5b), where the competition between the equally long vertical and horizontal boundaries is unresolved. Thus, we conclude that the boundary-cell model describes the experimental data much better than the inner-cell model (Fig. 4 and Extended Data Fig. 5).

**Figure 4.**
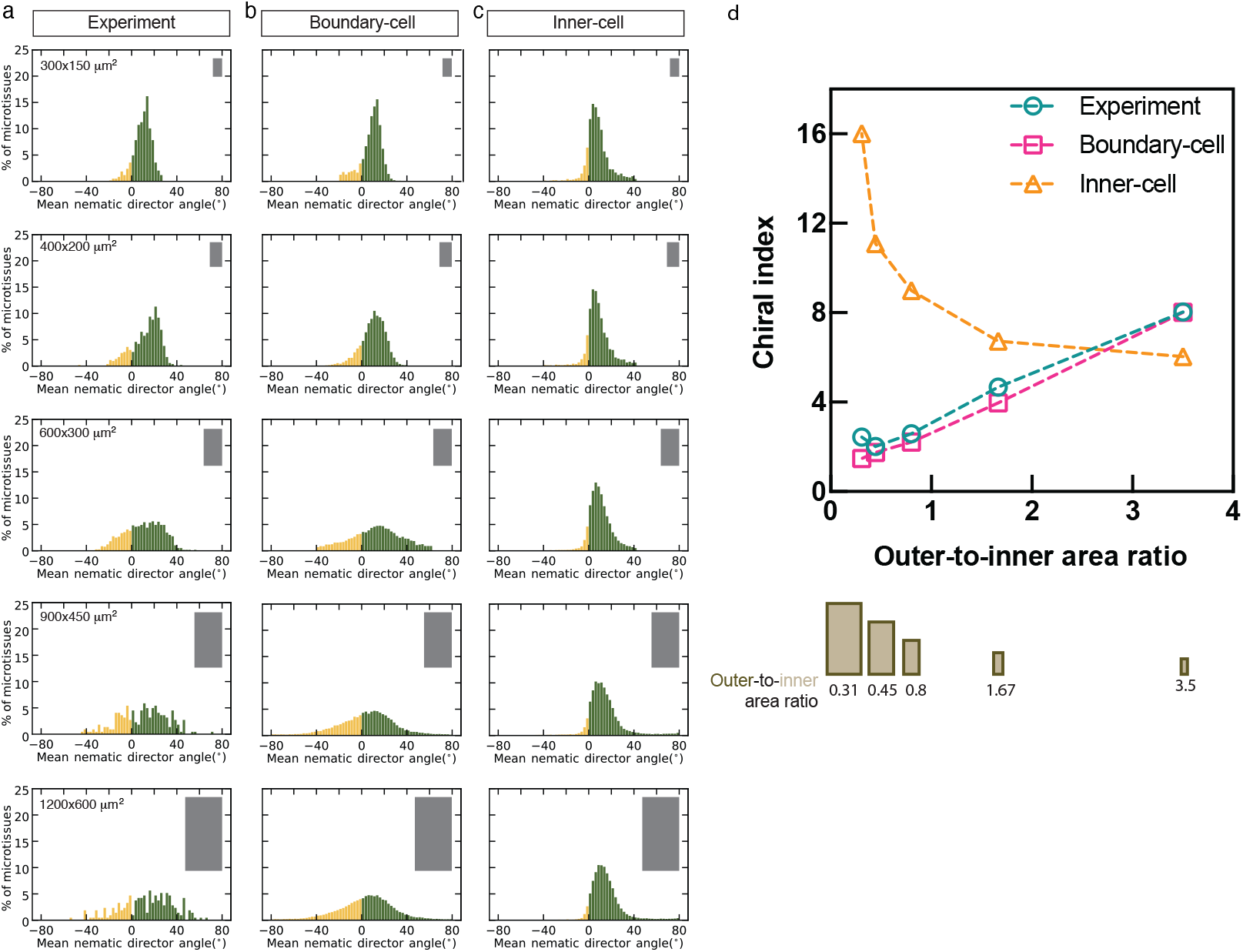
Boundary-cell model accounts for decreased microtissue chirality in larger areas. **a**,**b**,**c**, Histograms showing distributions of the mean nematic director angle for microtissues 48 h after seeding on various rectangular micropatterns of increasing area (from top to bottom) at a constant 1:2 aspect ratio. Experimental data (**a**) are compared with simulation outputs from the boundary-cell model (**b**) and the inner-cell model (**c**). The dimensions of the rectangular islands are indicated at the top left of the histograms in **a** and depicted as grey rectangles at the top right in **a** to **c**. The number of microtissues analysed is 587 (300×150 μm^2^), 522 (400×200 μm^2^), 579 (600×300 μm^2^), 223 (900×450 μm^2^), and 213 (1200×600 μm^2^) for experiment, and 100 simulations for each condition. Negative and positive values are coloured yellow and dark green bars respectively. **d**, Graph showing correlation of the chiral index versus the ratio of outer-to-inner area in experiments (teal circle symbol and dashed line) and the two models, boundary-cell model (magenta square symbol and dashed line) and the inner-cell model (yellow triangle symbol and dashed line). Calculated values of the area ratio are listed below each corresponding pattern. Dark brown and light brown represent outer and inner areas respectively. The outer area is defined as the 50 µm-wide zone adjacent to the micropattern boundary.

### Dynamics of establishing chiral cell alignment in rectangular 2D-microtissues

We further examined how chiral cell alignment developed in time. A series of live imaging experiments (Fig. 5a) revealed that direction of collective alignment (average island nematic director) emerged already in the first 6-12 hours (Fig. 5b,c) and remained steady up to 48 hours (Fig. 5a-c). During this period, the degree of alignment apparently increased (Fig. 5a) and the standard deviation between the values of nematic directors in different islands decreased (Fig. 5c). At the same time, measurement of local nematic directors revealed that the rate of establishing of chiral orientation is different in different regions of the islands (Fig. 5d). The nematic directors in the ‘vertical’ columns adjacent to the island long boundaries were uniformly tilted relatively to the boundary at already 3 hours post cell seeding and further only slightly decreased with time (Fig. 5e). In the columns distant from the boundaries, the nematic directors were initially characterised by high standard deviation which then gradually decreased with time, approaching the minimal value at ∼36-48 hours (Fig. 5e).

**Figure 5.**
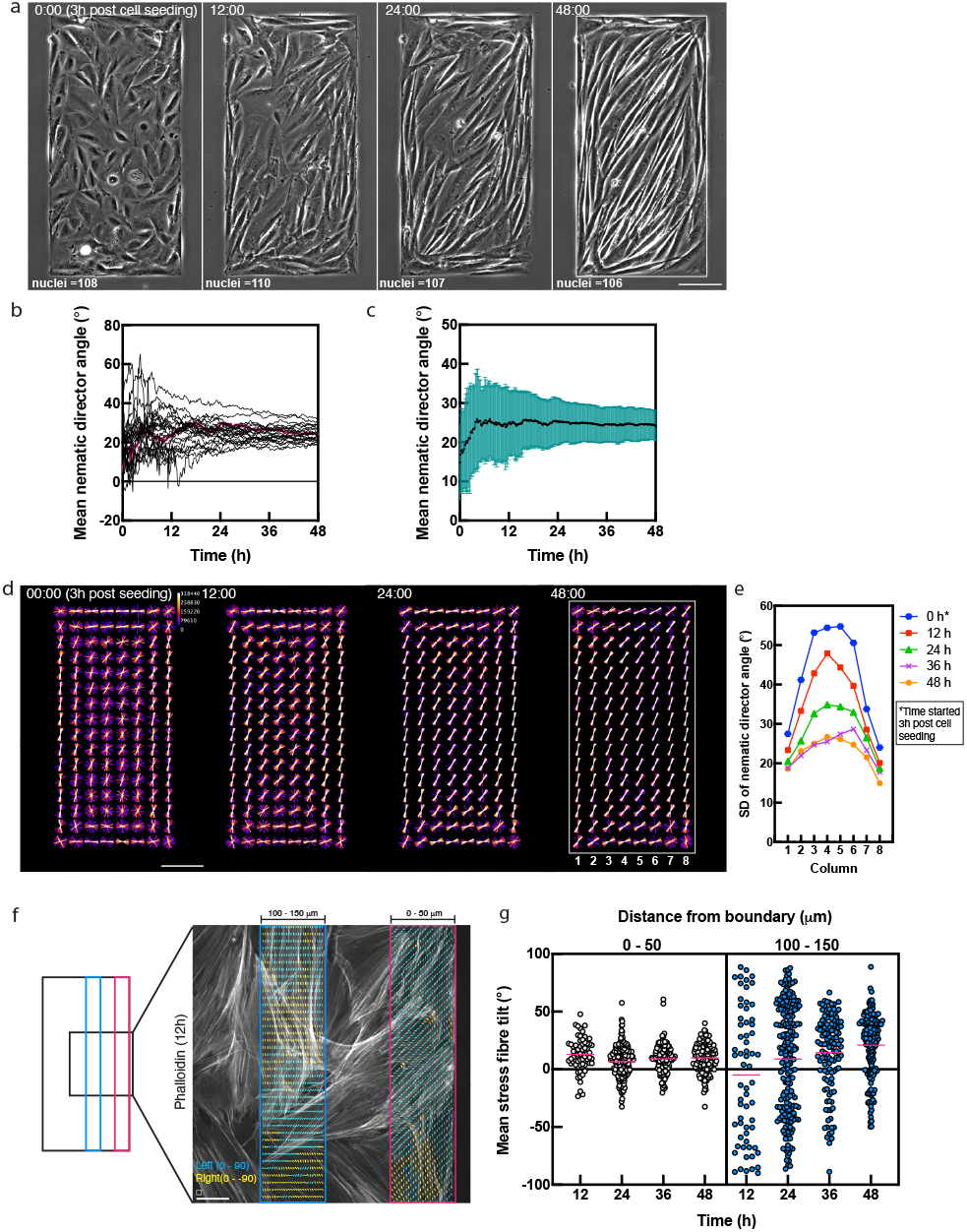
Outside-in propagation of chiral cell alignment and stress fibres orientation in rectangular 2D-microtissues. **a**, Time-lapse sequence of phase-contrast images showing a developing microtissue on a 300×600 μm^2^ rectangular micropattern. Timepoints (hh:mm) started 3 h post-cell seeding. The number of nuclei is indicated at each time point. Scale bar, 100 μm. **b**, Mean nematic director angles plotted over time for 23 individual microtissues. The red line represents the trajectory corresponding to the microtissue in **a**. **c**, Mean nematic director angle averaged across all 23 microtissues from **b**. The plot shows the mean (black line) ± s.d. (teal shading). **d**, Superimposition of nematic director fields from 23 microtissues at the indicated time points (hh:mm). The colour scale (Fire LUT; fluorescence intensity (a.u.) in a 32-bit range) represents the frequency of overlap. The microtissue is divided into eight columns (1 to 8) for the spatial analysis shown in **e**. Scale bar, 100 μm. **e**, Quantification of alignment variability, plotted as the standard deviation (SD) of the mean nematic director angle within each of the eight columns defined in **d**. *Time is shown in hours (h) from the start of imaging (3 h post-cell seeding). See also Extended Data Fig. 6. **f**, Analysis of stress fibre orientation in fixed microtissues. 2D-microtissues on 300×600 μm^2^ islands were fixed and labelled with phalloidin to visualise stress fibres at specific time points (hours, h) post-cell seeding. (Left) Diagram illustrating the imaging field of view (black square) relative to the microtissue (black rectangle). Stress fibre tilt was measured in a 50 µm-wide zone adjacent to the boundary (outer, magenta) and in a zone 100–150 µm from the boundary (inner, blue). (Right) Representative fluorescence image of phalloidin-labelled stress fibres in a microtissue fixed 12 h post-seeding. The local stress fibre tilt, measured in 20×20 pixel windows (box above scale bar), is depicted as a vector field overlay. Yellow and cyan vectors indicate left-tilted (–90° to <0°) and right-tilted (>0° to 90°) fibres respectively. For each image, the mean of all vectors in a zone is calculated and plotted as a single data point in **g**. Scale bar, 25 µm. **g**, Scatter plot of mean stress fibre tilt angle in the outer (0–50 µm zone; white dots) and inner (100–150 µm zone; blue dots) regions illustrated in **f** at different time points post-seeding. Each data point represents the mean angle from a single image, as described in **f**. Magenta lines indicate the mean value for each group. The number of images analysed (n) at each time point is: 63 (12 h), 213 (24 h), 124 (36 h), and 169 (48 h).

In addition to measurements of nematic directors in phase-contrast images, we performed similar measurement for the cells stained with phalloidin-AlexaFluor488 to reveal the orientation of stress fibres at different time points and different zones of the island (Fig. 5f). Consistently, these measurements showed that orientation of stress fibres in the columns adjacent to the vertical rectangular boundaries approached steady state in approximately 12 hour and characterised by relatively low standard deviation over time (Fig. 5g). At the same time, the standard deviation of stress fibres orientation in inner column (100-150 μm away from the boundary) was very high at 12 hours after seeding and gradually decreased with time (Fig. 5g). The average value of stress fibres orientation in these cells gradually increased and approached the maximum at 48 hours (Fig. 5g). Altogether, these measurements are in a good agreement with the boundary-cell model (which is illustrated by computational results shown in Extended Data Fig. 6), for the simple reason: in the model, the chiral order is informed by the tilted cells at the island boundaries, and, with time, propagates inward.

### Collective cell alignment in the mirror-symmetrical adhesive islands

In experiments described above, we investigated the alignment of cells in rectangular 2D-microtissues. Rectangle is a non-chiral figure (it has left-right symmetry) and was used to draw out collective effects of the intrinsic chirality of the cells. In this section, we examined how the chiral cells aligned inside the islands with the chiral shape. The non-rectangular parallelograms are chiral in the sense that they cannot be superimposed with their mirror image (Fig. 6). Thus, we used the parallelograms preserving the area and ratio between the short and long sides equal to our standard 300×600 μm^2^ rectangle but having 60° and 120° tilt angle instead of 90° (Fig. 6a).

**Figure 6.**
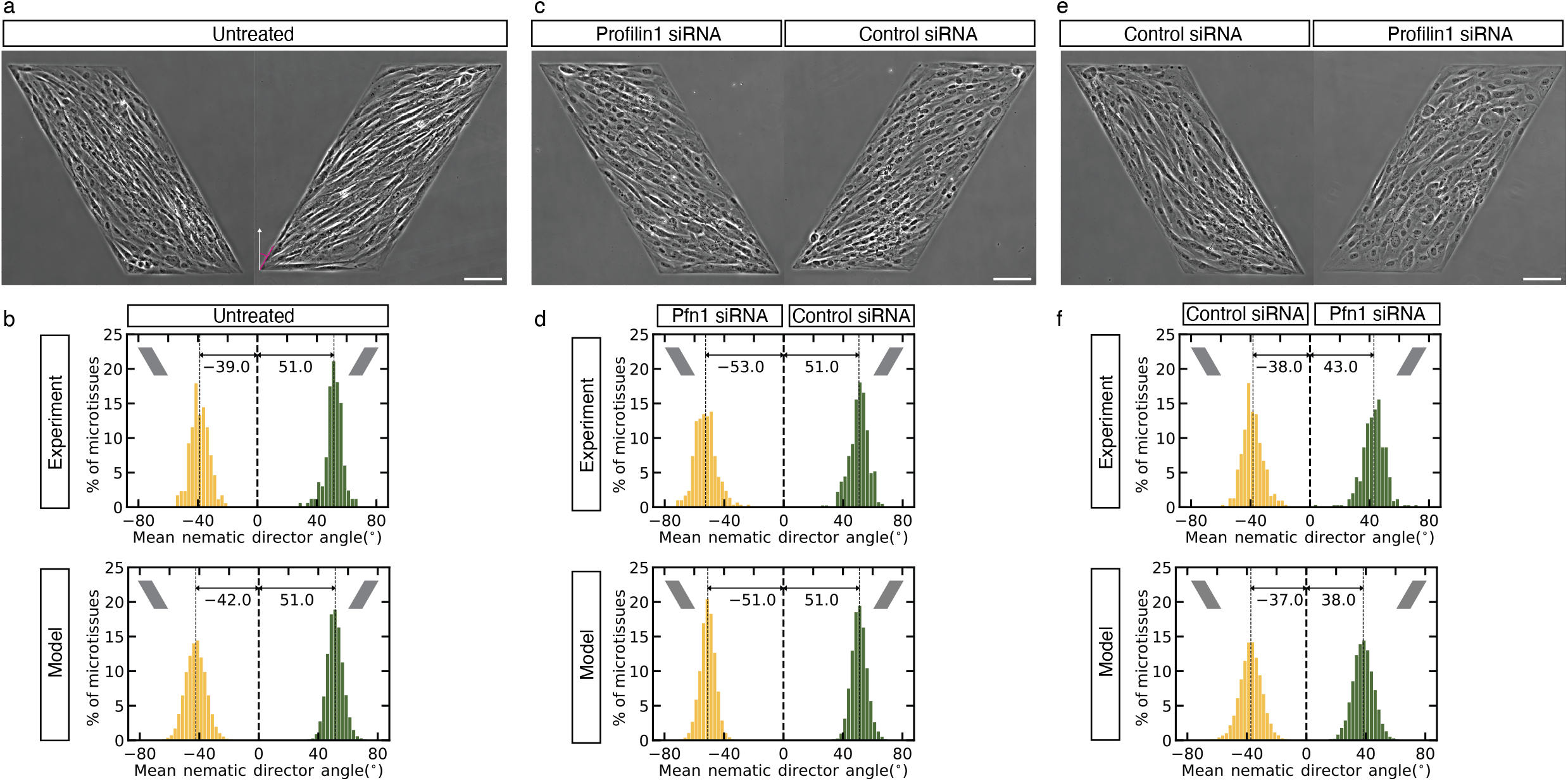
Collective chiral cell alignment on mirror-symmetrical micropatterns. **a, c, e**, Phase-contrast images of cells 48 hours after plating on mirror symmetrical left-tilted or right-tilted parallelogram micropatterns. Cells were either untreated, transfected with control siRNA, or transfected with profilin-1 siRNA as indicated. The magenta line in **a** shows a 30° angle relative to the perpendicular from the parallelogram base (white arrow). This perpendicular defines the 0° reference for all angle measurements depicted in the histograms. Scale bars, 100 µm. **b, d, f**, Histograms showing distributions of mean nematic director angles characterising individual microtissues under the indicated conditions 48 h after plating in experiment and outputs from boundary-cell model simulations. The number of microtissues (n) analysed for experiments were: (**b**) 166 (right-tilted) and 173 (left-tilted) for untreated cells; (**d**) 406 (control siRNA, right-tilted) and 297 (profilin-1 siRNA, left-tilted); and (**f**) 347 (profilin-1 siRNA, right-tilted) and 312 (control siRNA, left-tilted). For simulations, n = 100 for each condition.

Measurement of nematic directors of the 2D-microtissues confined to these chiral patterns revealed that average orientations of cells in these micropatterns were not mirror-symmetrical. While in the right-tilted parallelogram the average nematic director was oriented at 51° relative to the vertical axis, the nematic director in left-tilted parallelogram was significantly different – not -51° but -39° (Fig. 6b). These nematic director values were faithfully reproduced by computations based on boundary-cell model (Fig. 6b). Qualitatively, the difference is easy to understand: in the right-tilted parallelogram, the right tilt of the boundary cell synergises with the right tilt of the island, while in the left-tilted parallelogram, the right tilt of the boundary cell competes with the left tilt of the island.

Thus, the intrinsic chirality of cells interacts with the chirality of the island shape and converts the mirror-symmetrical parallelograms into non-symmetrical microtissues. The question then emerged whether mirror symmetrical micro-tissues could be created inside the islands with mirror symmetrical shapes. From our previous work^23^, we know that some genetic perturbations can reverse the direction of individual cell chirality which also reversed direction of collective cell orientation in rectangular islands. We choose one of these perturbations, depletion of profilin 1, and compared the alignment of profilin-1 knockdown and control cells confined to the chiral parallelogram islands (Fig. 6c,d,e,f). These experiments showed that profilin1-depleted cells exhibited mirror-reflected alignment of the control cells when they are confined to the mirror-reflected parallelograms (Fig. 6c,e). These experimental measurements were also faithfully reproduced by computations based on boundary-cell model (Fig. 6d,f). These results are easy to understand as follows: the left-tilted profilin-1 knockdown cells in the left-tilted parallelogram generate the collective alignment axis tilted to the left by approximately 50° (Fig. 6c,d), same tilt but to the right generated by the right-tilted control cells in the right-tilted parallelogram (Fig. 6a,b,c,d). Similarly, the left-tilted profilin-1 knockdown cells in the right-tilted parallelogram generate the diminished collective tilt to the right by approximately 40° (Fig. 6e,f), same tilt but to the left generated by the right-tilted control cells in the left-tilted parallelogram (Fig. 6a,b,e,f). Interestingly, increase of the parallelogram size reduced the deviation from mirror symmetry in the collective cell orientation emerging due to chiral interaction of the cell with the boundary (Extended Data Fig. 7). In the similar parallelograms with 2-fold increase in the perimeter, the simulation predicted that both control and profilin-1 knockdown cells will demonstrate symmetric average tilt following the direction of the longer diagonal of the parallelograms (Extended Data Fig. 7).

## Discussion

Our study uncovers the mechanism of collective cell chirality emergence from interactions of a subcellular chiral behaviour of stress fibres with extracellular geometry. Namely, our findings support the boundary-cell model: cells near the adhesive island boundaries align with the boundaries but develop tilted stress fibre orientation and shape relative to the boundary. Then, mutual cell-cell alignment propagates inward into the bulk of the cell microtissues. This model correctly explains the nontrivial frequencies of several patterns on rectangular adhesive domains as the result of competition between opposing influences of perpendicular sides of the rectangle islands. We found that microtissue size and geometry significantly modulate chiral strength, with boundary effects dominating small systems and diminishing in larger ones.

The boundary-cell model further makes a nontrivial prediction about different alignment angles on mirrored parallelogram patterns, for both control and reverse-chirality cells, which is confirmed by the experiment. These experiments provide a first attempt to investigate (in a very simplified form) a general question on how the mirror symmetrical tissues and organs can be built of cells with intrinsic left-right asymmetry. The theory shows that formation of mirror symmetrical small cell groups requires the switch of chirality of individual cells, while increase of the group size can diminish the role of sign of individual cell chirality. It is interesting whether any of these solutions are utilised in *in vivo* developmental processes.

The boundary-cell model correctly and quantitatively captures these features of multicellular patterns on larger adhesive domains, while the initially plausible inner-cell model fails to explain the data. We cannot exclude, however, that some elements of the inner-cell model can work synergistically *in combination* with the boundary-cell mechanism – future research needs to establish if this is the case.

The boundary-cell mechanism analysed in this study is not the first attempt to explain the formation of left-right asymmetric structures in 2D cell collectives. Several recent studies, both experimental and theoretical, have put forward other mechanisms. For example, examination of asymmetrical flows of cell monolayers on adhesive strips (up along one boundary of the strip vs down along the opposite boundary) revealed chiral rotating rows of multicellular vortices positioned at certain distance from the boundaries^49^. These rotations could induce the left-right asymmetry of the cell gliding directions^40, 49^. Advanced model of multicellular interactions based on hydrodynamic-like active polar gel theory suggests that chiral stresses in nematic cell monolayers induce collective chirality^42^. Several experiments and models suggest that topological defects in multicellular alignment maps play a crucial role in collective cell patterns^50, 51^, and some studies put these topological defects forward as principal drivers of the microtissue’s chirality^42, 49^. In our study, for the first time, the rules of chiral behaviour of individual cells were not proposed *ad hoc* but inferred from the careful examining of self-organisation of the actin cytoskeleton in the cells of different shapes. Novel methodological approaches suggesting a general way to inferring individual cell chiral behaviours and intercellular interaction rules from multicellular chiral patterns^52^ could open an avenue to cracking the chirality propagation puzzle in the future.

The greatest remaining puzzle concerns microscopic mechanisms behind the uncovered pathway of chirality propagation across the cellular scales. First part of this puzzle is the self-organisation of the collective rotational bias of cytoskeletal elements in single adhesive cells. Given the mechanochemical nature of cytoskeletal dynamics at large, it is very suggestive that a chiral mechanical torque rotates the stress fibres anchored at the focal adhesions at the cell periphery. This torque could stem from the hypothesised cooperative contraction of circumferential contractile bundles interacting with tilts of the radial stress fibres and biasing these tilts, assisted by formin-generated rotation of individual actin filaments in stress fibres^22^. Indeed the role of formins as well as several other actin-associated proteins regulating actin polymerisation and crosslinking in cell and organismal chirality is supported by numerous experimental studies ^23, 28-31, 35, 53-55^. However, intracellular chiral torque could also be driven by the interaction of actin filaments with some molecular motors including myosin II ^34, 37, 38^, myosin I^24-27,39^ and perhaps other myosins^36, 56^. While in all these scenarios the major asymmetric element was the helical polar actin filament, the methods of translating this asymmetry into larger scale cellular asymmetry could probably differ in different cellular types. It is especially interesting that some experimental perturbations of actin-associated proteins not simply destroy the cell chirality but reversed its sign suggesting that cells can exist in two mirror configurations (refs^23, 24, 27^ and present study). The exact nature of these subcellular mechanochemical torque generators and their reversion await further investigation.

Second, the nature of the intercellular propagation of the chiral sub-cellular architecture is yet to be understood. It could depend on the simple steric interaction and packing of cells elongated along the stress fibres bundles direction in dense cell groups, but also specific molecular pathways regulating cell-cell contacts as well as cell-matrix production and remodelling may be involved. The literature on the mechanism of cell alignment is abundant (see for example^57, 58^) but in this study we did not go deeply into any particular mechanism using just a simple phenomenological rule. Also, relative roles of the cell dynamic shapes, intracellular stress fibre alignment processes and intercellular interactions between these factors must be clarified.

Last but not least, the remaining challenge is understanding chirality mechanisms and their physiological significance in 3D, as physiologically relevant tissue dynamics often play out in 3D extracellular matrix (cells do exhibit chiral behaviour in 3D^54, 59^). Despite these gaps in knowledge, our findings provide a necessary step for understanding how chirality at the single-cell level scales to collective organisation and may offer insights into biological processes where cell chirality influences tissue morphogenesis.

## Acknowledgments

We thank R. Wedlich-Soldner (University of Münster, Germany) for LifeAct-GFP and Lucie S. E. Kim (MBI, Singapore) for expert help with illustration and the SIMBA microscopy facility and nanofabrication core facility at the Mechanobiology Institute for technical help. The research is supported by the Ministry of Education under the Research Centres of Excellence programme through the Mechanobiology Institute at National University of Singapore (grant numbers A-0003467-01-00 and A-0003467-00-00) and the National Research Foundation (NRF) Singapore, National University of Singapore under its Mid-Sized Grant (grant number NRF-MSG-2023-0001). The research is supported in part by the National Science Foundation (Grant DMS1953430 to A.M.).

## Author Contributions

W.S., A.D.B., A.M. and Y.H.T. conceived and designed the experiments and wrote the manuscript with input from all the authors. D.T.L.N. and Y.H.T. performed most experiments. W.J.G. and R.B.T. contributed to some experiments. W.J.G., H.T.O., C.F., and Y.H.T. performed image and data analysis. W.S. and A.M. developed the computational model and performed all simulations.

## Competing Interests

The authors declare no competing interests.

## Additional Information

Supplementary Information includes 7 Extended Data Figures, and 3 supplementary videos can be found with this article.

## Data Availability

All data generated or analysed during this study are included in this published article (and its Supplementary Information files). Raw datasets corresponding to any graphs presented in this study are available from the corresponding authors on reasonable request.

## Code Availability

Custom-written codes used to analyse the data in the current study are available on Zenodo.org, https://zenodo.org/records/17296461.

## Extended Data Figure Legends

**Extended Data Fig. 1.**
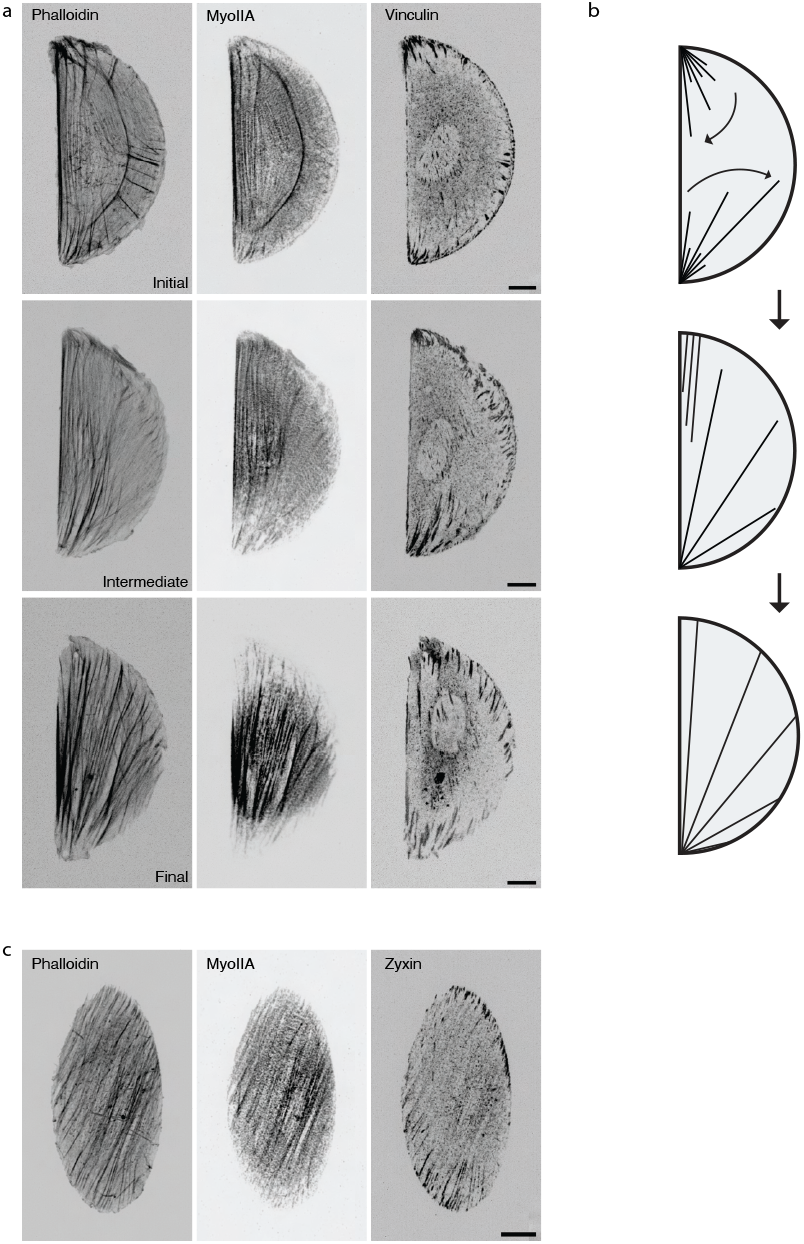
Actin stress fibres and focal adhesion organisation in individual confined cells. **a**, Examples of stress fibre organisation and focal adhesion distribution in cells confined to a semi-circular micropattern 6 h after plating. The images display the proposed temporal stages of cytoskeletal evolution – “initial,” “intermediate,” and “final”, which are illustrated schematically in **b**. **b**, Schematic illustrating the proposed progression of actin cytoskeleton self-organisation, from initial radial fibres formation and subsequent fibres tilting to the emergence of uniformly right-tilted stress fibres relative to the semi-circular diameter. **c**, Example of stress fibre organisation in a cell confined to an elliptical micropattern. Note that the stress fibres are tilted to the right relative to the major axis of the ellipse, as previously reported in Tee *et al*. (2023). In **a** and **c**, actin and myosin IIA were visualised with phalloidin and an anti-myosin IIA antibody, respectively. Focal adhesions were visualised with an anti-vinculin antibody in **a** and an anti-zyxin antibody in **c**. Scale bars, 10 µm.

**Extended Data Fig. 2.**
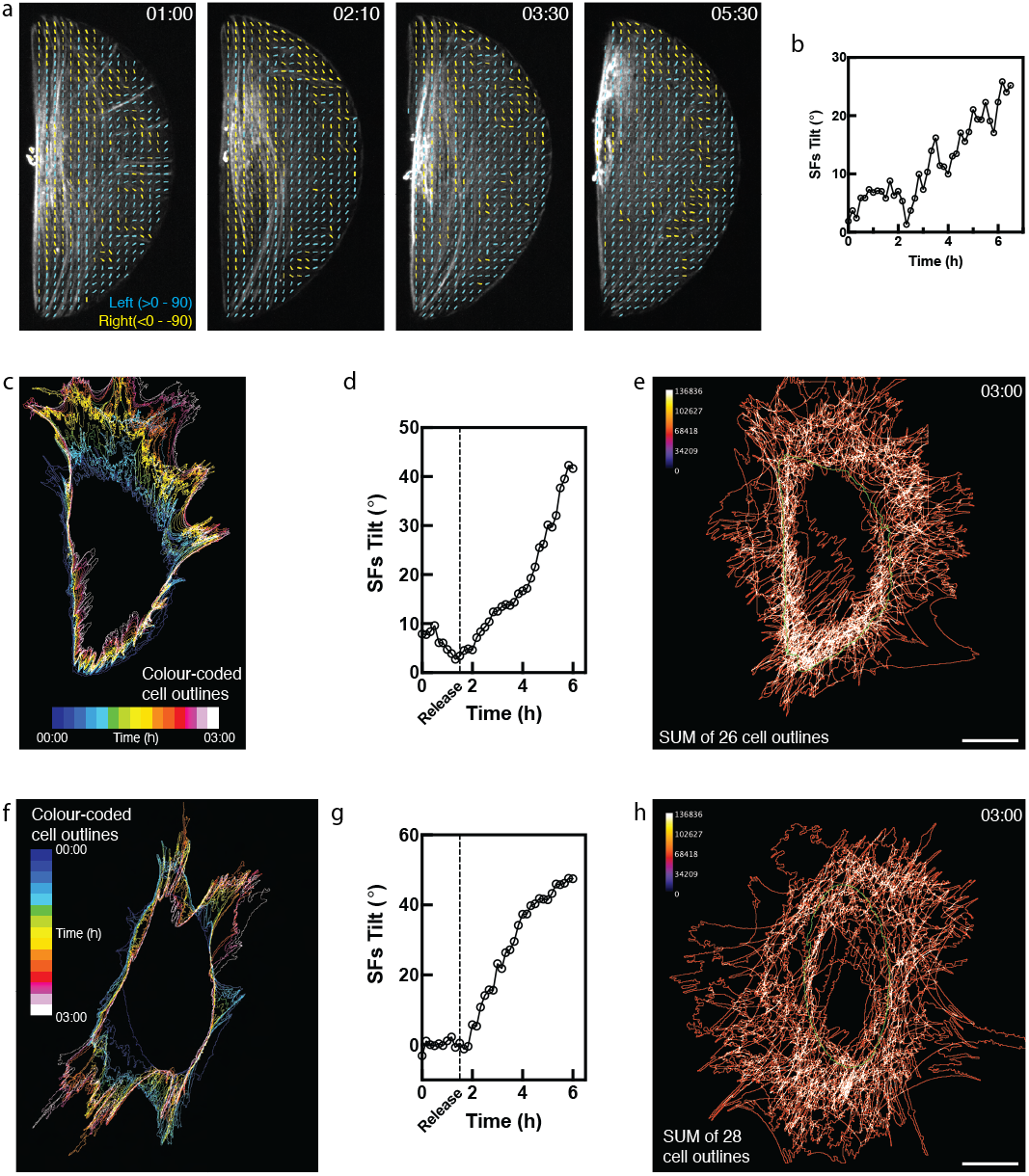
Actin fibre tilt and cell deformation following cell release from confinement. **a**, Vector fields of local actin alignment based on SpyFastAct-labelled actin cytoskeleton at selected time points (hh:mm) for the cell shown in Fig. **1c**. Vectors are coloured by their tilt direction relative to the vertical diameter of the semi-circle (0°): yellow for left-tilted (–90° to <0°) and cyan for right-tilted (>0° to 90°). The circular mean of all vectors was calculated for each time point and plotted in **b**. **b**, Stress fibres (SFs) tilt angle over time (in hours, h) for the cell shown in **a. c, f**, Temporal colour-coded overlays of the cells’ outlines shown in Fig. **1e** and **1g**, respectively. **d, g**, Quantification of the mean SFs tilt over time for the cells shown in Fig. **1e** and **1g**. Time (in hours; h) begins at the start of imaging; release from confinement occurred at t = 1.5 h. **e, h**, Density maps generated from superimposed outlines of cells 3 h after release from semi-circular (n = 26, **e**) or elliptical (n = 28, **h**) confinement. The colour scale (fluorescence intensity (a.u.) in a 32-bit range) represents the frequency of overlap, indicating the probability density of the cell boundary. Scale bars, 20 µm.

**Extended Data Fig. 3.**
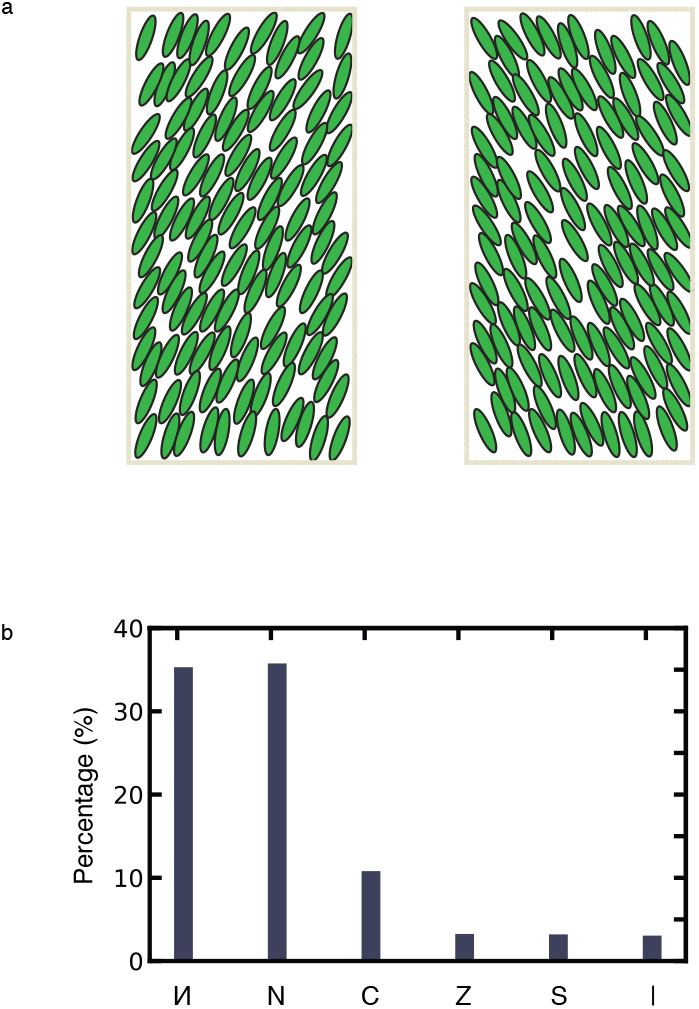
Predicted patterns of alignment of non-chiral cells in rectangular microtissues. **a**, Simulation outputs showing the emergence of the (left) “И”- and (right) “N”-orientation patterns. Individual cells are depicted as green ellipses and the micropattern boundary is outlined in light brown. **b**, Frequency distribution of the six orientation patterns predicted by simulation of non-chiral cells. Note that orientation patterns “И” and its mirror-image “N” occurs with equal frequency. See also Extended Data Fig. 4.

**Extended Data Fig. 4.**
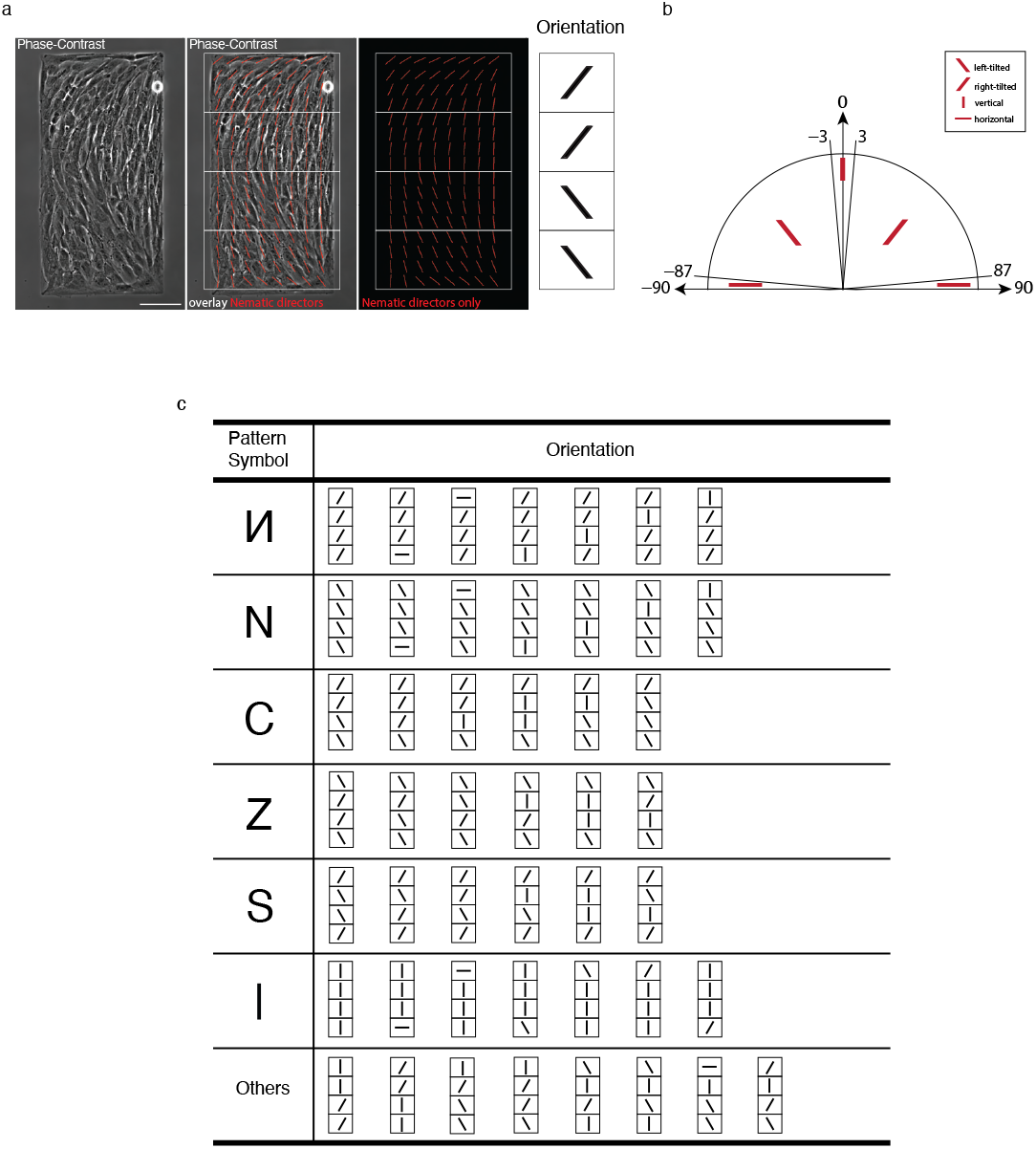
Types of collective cell alignment patterns in rectangular microtissues. **a**, Method for classifying microtissue orientation patterns. (Left) A phase-contrast image of cells on a 300×600 µm rectangular micropattern 48 h after plating. (Middle) The same image overlay with the local cell orientation field (nematic directors, red lines). (Right) To classify the overall orientation pattern, the microtissue is divided into four quadrants (white boxes). The mean nematic director angle is calculated for each quadrant and assigned a symbol (/, \, |, or –) according to the criteria defined in **b**. See also **Fig. 3a**. **b**, Definition of the qualitative orientation symbols. The panel shows the range of values of mean nematic director angles used to classify each quadrant as right-tilted (/), left-tilted (\), vertical (|), or horizontal (–). **c**, Table showing the combinations of quadrant symbols that define the six primary orientation patterns. The ‘Others’ category comprises other observed combinations that do not fit in any of these classes, representing approximately 2.5% of the total population. See Methods for further details on the classification.

**Extended Data Fig. 5.**
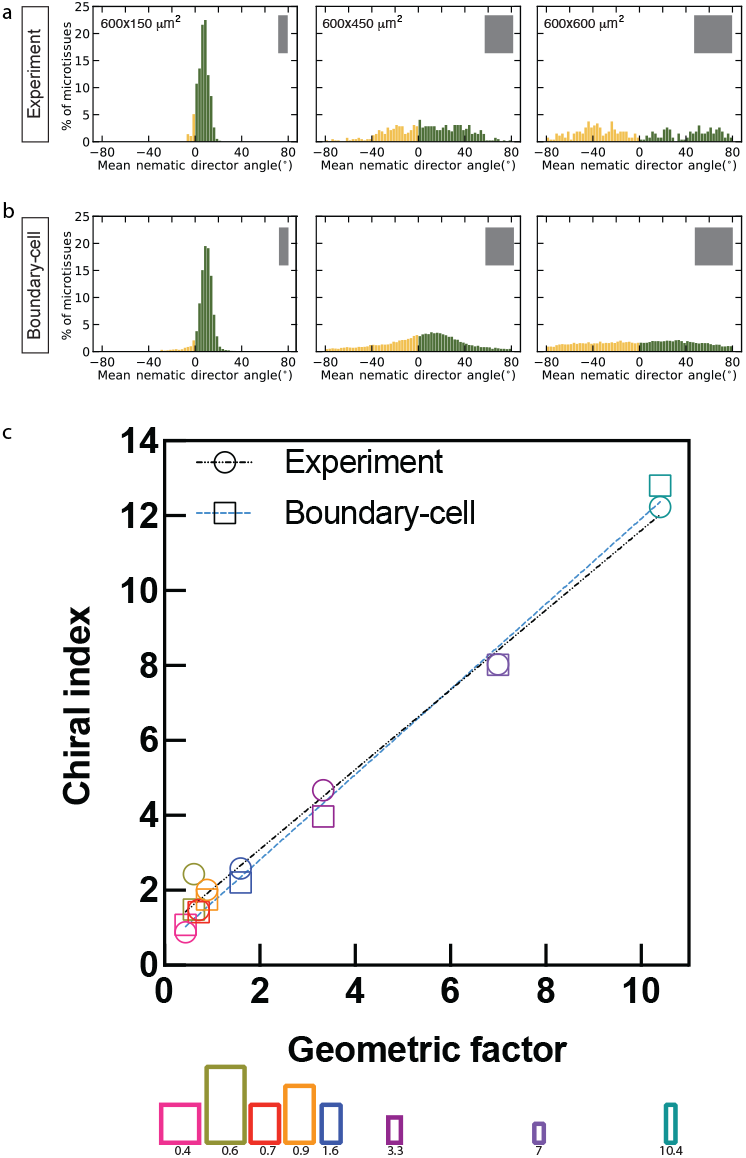
Chirality of cell alignment in microtissue as a function of microtissue area and aspect ratio. **a, b**, Comparison of mean nematic director angle distributions at 48 hours for experimental microtissues (**a**) and corresponding boundary-cell model (**b**). The dimensions of the rectangular islands are indicated at the top left of the histograms in **a** and depicted as grey rectangles at the top right in **a** and **b**. The number of microtissues analysed is 569 (600×150 μm^2^), 426 (600×450 μm^2^), and 293 (600×600 μm^2^) for experiment, and 100 simulations for each condition. Negative and positive values are coloured yellow and dark green respectively. **c**, Correlation between the chiral index and a geometric factor of experimental data (circle and dotted black line) and boundary cell simulation results (square and dotted blue line). The geometric factor is defined as the product of the rectangular island’s aspect ratio and the ratio of its outer-to-inner area. Values for the geometric factor are listed below its corresponding micropattern. Dotted lines represent linear best-fits.

**Extended Data Fig. 6.**
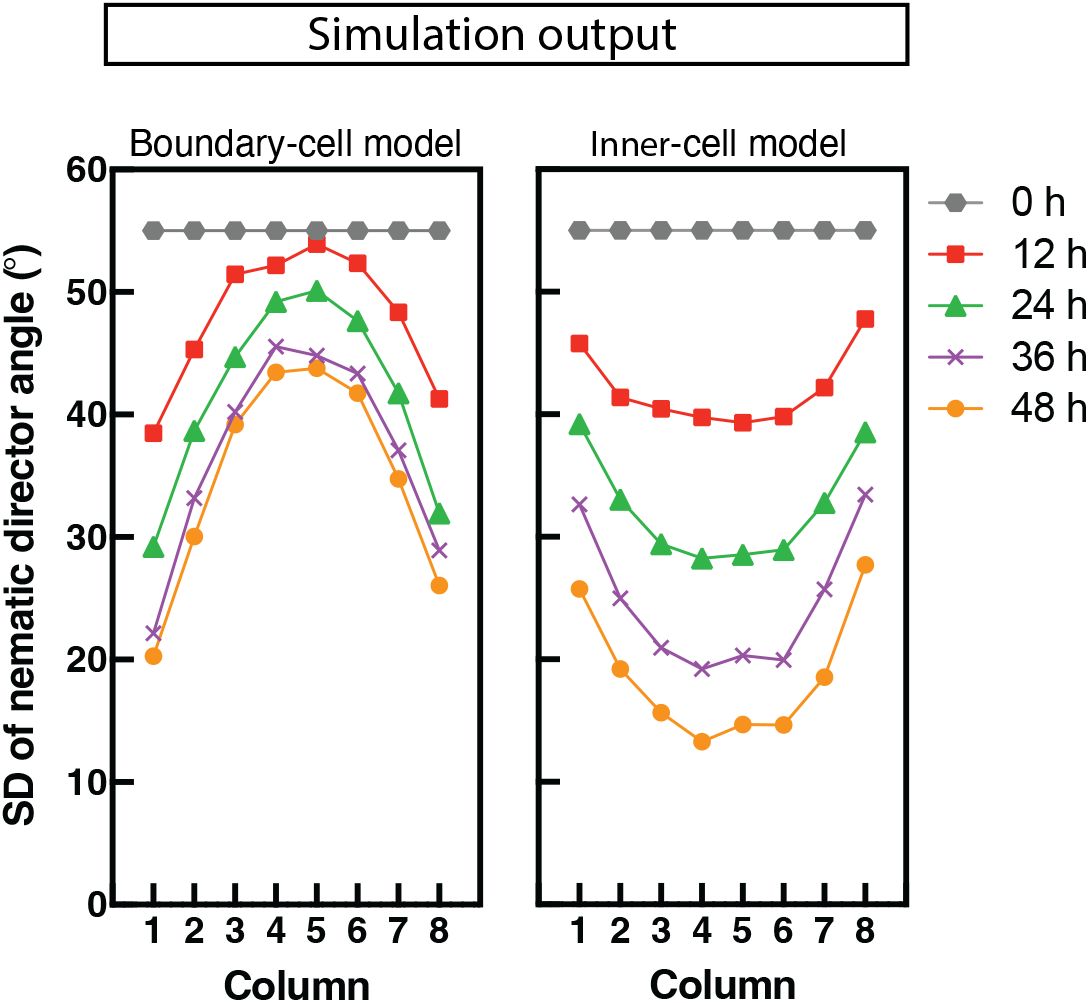
Temporal evolution of cell alignment variability as predicted by two distinct models. The plots show the standard deviation (s.d.) of the mean nematic director angle within each of the eight columns across the microtissue as determined by simulations. Both simulations were initialised with a uniform, high s.d. across all columns at t = 0 h of modelling. (Left) In the boundary-cell model, the s.d. of cells near the boundary decreases faster than that for the inner cells until reaching steady states where the outer cells are much more ordered than the inner cells, in line with the experiments. See also Fig. **5e**. (Right) The inner-cell model predicts the opposite trend, with the s.d. of the outer cells decreases slower than that for the inner cells. This outcome is inconsistent with experimental observations.

**Extended Data Fig. 7.**
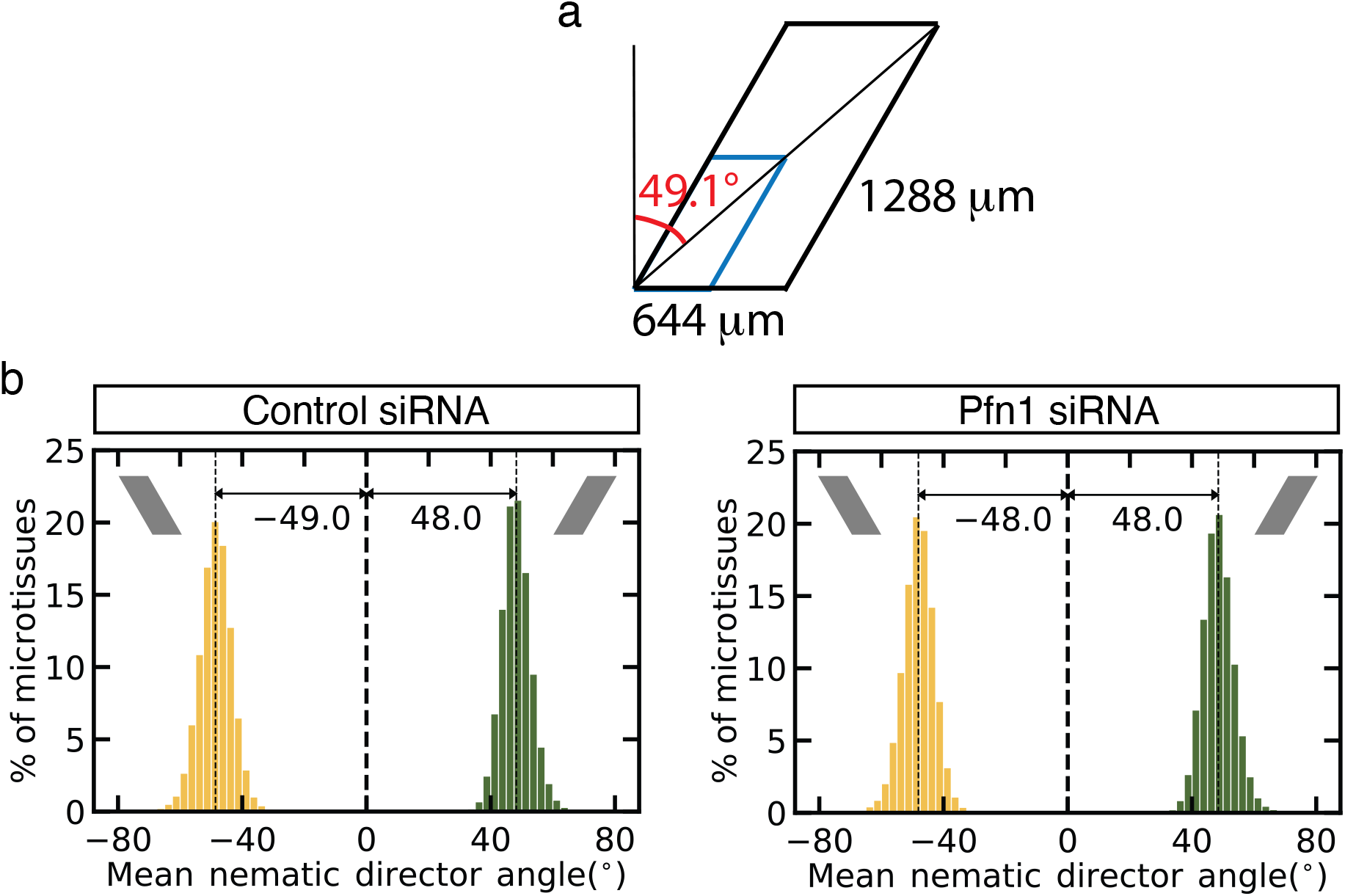
Increase of size of mirror symmetrical micropattern abolishes the effect of individual cell chirality on collective cell orientation. **a**, Schematic illustrating the geometry of the right-tilted parallelogram micropatterns. The calculated angle between the longer diagonal and the axis perpendicular to the base is 49.1° (red). For reference, the smaller micropattern used in the experiments in Fig. 6 (322 µm × 644 µm) are outlined in blue. **b**, Histogram showing the distribution of mean nematic director angles from boundary-cell model simulations (n = 100) of control siRNA-treated or profilin-1 siRNA treated cells on the larger mirror symmetric parallelogram micropatterns (644 µm × 1288 µm). The angles are measured relative to the perpendicular from the parallelogram’s base, and the mean of the distribution is indicated.

## Supplementary Video Captions

**Supplementary Video 1**| Self-organisation of the actin cytoskeleton in single cell plated on a semi-circular fibronectin island. (Left) Fluorescence images of actin (green, SpyFastAct) and the nucleus (red, Hoechst-33342) over time (hh:mm). (Right) Corresponding vector field of local actin alignment. Vectors are coloured-coded: yellow for left-tilted (–90° to <0°) and cyan for right-tilted (>0° to 90°). Imaging began 2 h post-seeding. Images were recorded at 10-minute intervals over a period of 6.5 hours. Display rate is 10 frames/sec and corresponds to the time-lapse series in Fig. 1c and Extended Data Fig. 2a.

**Supplementary Video 2**| Actin cytoskeleton and cell dynamics following release from semi-circular island confinement. Actin was visualised with SpyFastAct-650. Release was induced by adding streptavidin-fibronectin (SA-FN) to PLL-PEG-biotin passivated surface at the 01:30 (hh:mm) mark of the video. Imaging began 6 h post-seeding and was recorded at 10-minute interval for ∼5 hours. Display rate is 10 frames/sec and corresponds to the time-lapse series in Fig. 1e.

**Supplementary Video 3**| Actin cytoskeleton and cell dynamics following release from elliptical island confinement. Actin was visualised with SpyFastAct-650. The video begins at the moment of release (time 00:00), which was induced by adding streptavidin-fibronectin (SA-FN) to the passivated surface. Frames were captured at 10-minute intervals for ∼5 hours. Display rate is 10 frames/sec and corresponds to the time-lapse series in Fig. 1g.

## Methods

### Cells and siRNA transfection

Human foreskin fibroblasts (HFF) from American Type Culture Collection (catalogue no. SCRC-1041) were cultured in Dulbecco’s modified Eagle’s medium high glucose supplemented with 10% fetal bovine serum (FBS), 1 mM sodium pyruvate and antibiotics (10 Units/mL penicillin and 10 Units/mL streptomycin) at 5% CO_2_ at 37°C. Cells used for experiment were between passage number 5 to 15. For siRNA transfection, cells were seeded into a 35 mm dish on day 0 and transfected with 100 μM of siRNAs using Lipofectamine RNAiMAX (catalogue no. 13778150) on days 1 and 2. siRNA-transfected cells were trypsinised on day 3 and replated onto micropatterned substrates. siRNAs used in this study were control siRNA (Dharmacon, ON-TARGETplus Non-targeting control, catalogue no. D-001810-01) and profilin 1 siRNAs (target sequences, 5’-GCAAAGACCGGUCAAGUUU-3’ and 5’-CACGGUGGUUUGAUCAACA-3’; RNAs synthesised by GenScript Biotech (Singapore) Pte. Ltd). All cell culture and transfection reagents were obtained from Invitrogen. Other chemicals and reagents were obtained from Sigma, unless otherwise stated.

### Micropatterning of substrates

Micropatterns used for the individual cell experiment include: circular micropattern (48 µm-diameter), elliptical micropattern of 1:2 aspect ratio (34:68 µm) and semi-circular micropattern (80 µm-diameter). Micropatterns used for the 2D-microtissue experiment include: rectangular micropatterns (150×300 µm; 200×400 µm; 300×600 µm; 450×900 µm; 600×1200 µm; 150×600 µm; 450×600 µm and 600×600 µm) and mirror symmetrical parallelograms (base: 322 µm, long side: 644 µm, ±30° tilt from perpendicular from base). Each micropatterned substrate was fabricated by stencil patterning as previously described in our earlier work^23^. Briefly, polydimethylsiloxane (PDMS) (Sylgard 184 kit, Dow Corning) was cast on the photoresist mould, containing micropattern designs of interest, using a 10:1 ratio (w/w) of elastomer to crosslinker and cured for 2 h at 80°C. The crosslinked PDMS layer was peeled off and stamps were cut out manually. The PDMS stamp was then inverted and placed onto a hydrophobic uncoated 35 mm µ-dish (ibidi GmbH; catalogue no. 80131). Norland Optical Adhesive 73 (NOA-73, Norland Inc.) was deposited along an edge of the stamp and allowed to flow through the gaps between the PDMS stamp and dish by capillary action, upon which the stamp was sealed on all sides using NOA-73. The NOA-73 stencil was cured under ultraviolet illumination using 50% power for 13 seconds (UV-KUB UV-LED, Kloè). After peeling off the PDMS stamp, the stencil and dish were incubated with fibronectin (Calbiochem, Merck Millipore) at a concentration of 50 µg ml^−1^ in 1×PBS at 4°C overnight after a brief degassing at 10 mbar. At the end of the incubation, the fibronectin solution was aspirated, and the stencil was removed. The printed dish bottom was passivated with 0.2% Pluronic acid-H_2_O for 10 min. Finally, the passivated dishes were washed thrice with 1×PBS before cell seeding.

### Dynamic micropatterned substrate

Micropatterned substrate was fabricated by stencil patterning as described above with the following modification. After fibronectin incubation, and the stencil was removed, the printed dish was passivated with 0.1 mg ml^−1^ of PLL(20)-g[3.5]-PEG(2)/PEG(3.4)-biotin(50%) (SuSoS AG, Switzerland) either overnight at 4°C or for 1 hour at 37°C. Finally, the passivated dishes were washed thrice with 1×PBS before cell seeding. To release cells from confinement, streptavidin-conjugated fibronectin (Streptavidin Conjugation Kit - Lightning-Link®, Abcam, catalogue no. ab102921) were added to the medium during imaging to a final working concentration of 750 ng cm^-2^.

### Labelling of actin and nuclei for live cell imaging

To visualise actin cytoskeleton in cells during live imaging, cells were either transfected with LifeAct-GFP DNA plasmid via electroporation (Neon® transfection system, Life Technologies) following manufacturer’s instructions, or labelled using SPY650-FastAct™ (Cytoskeleton Inc., catalogue no. CY-SC505). Electroporation condition consists of two pulses of 1150 V for 30 milliseconds. FastAct probe was used at 1:2000 dilution and cells were labelled for at least 2 hr before proceeding to imaging. FastAct probe remained in the medium during imaging. For living imaging of nuclei, nuclei were labelled with Hoechst 33342 (10 μg ml^−1^ for 10 min).

### Immunofluorescence antibody staining

Cells were seeded on printed dishes containing circular, elliptical, or semi-circular micropatterns at a density of 5×10^4^ cells ml^-1^ for 10 min. The medium containing unattached cells was then replaced with fresh DMEM. After 6 h incubation, the cells were fixed using 4% paraformaldehyde (Tousimis, USA) in PBS for 10 min, followed by three 1×PBS washes. Cells were permeabilised using 0.5% Triton-X-100 in PBS, and then blocked with 5% bovine serum albumin (BSA)-PBS for 1 h at room temperature (RT) before incubation with appropriate primary antibodies overnight at 4°C: anti-vinculin (Sigma, catalogue no. V9131, dilution 1:300, clone hVIN-1); anti-myosin IIA (Sigma, catalogue no. M8064, dilution 1:1000) and anti-zyxin (Santa Cruz, catalogue no. sc-293448, dilution 1:300). Cells were then incubated for 45 min at room temperature with appropriate AlexaFluor-conjugated secondary antibodies (Molecular Probes, dilution 1:500). Actin and nucleus staining were performed using phalloidin (Molecular Probes) and Hoechst-33342 (Invitrogen) respectively.

### Live cell imaging and confocal microscopy

For live cell imaging experiment, cells were seeded on micropatterns at a density of 5×10^4^ cells ml^-1^ for 10 min. The medium containing unattached cells was then replaced with Leibovitz’s L-15 containing 10% FBS. Time-lapse images for 5-10 h at 10 min intervals and Z-stacks of step-size 0.35 μm with total height of 10-12 µm were acquired with a spinning disc confocal microscope (PerkinElmer Ultraview VoX) attached to an Olympus IX81 inverted microscope, equipped with a 60× oil immersion objective (1.35 NA, UPlanSApo), an EMCCD camera (C9100-13, Hamamatsu Photonics) for image acquisition, and Volocity software (PerkinElmer) to control the set-up. Imaging of cells on dynamic micropatterned substrate was performed on spinning disc confocal microscopy coupled with the live super-resolution (SR) module (Roper Scientific) attached to a Nikon Eclipse Ti-E inverted microscope with Perfect Focus System, equipped with 100× oil immersion objective (1.4 NA, PL APO VC), a sCMOS camera (Photometrics Prime 95B) for image acquisition, and MetaMorph software (Molecular Devices) to control the set-up. Fixed samples were also imaged with the above step-up using a 100× oil immersion objective (1.4 NA, PL APO VC), or on the BC43 benchtop spinning disc confocal (Andor) using a 60× oil immersion objective (1.40 NA, UPlanSApo). Maximum projection of the Z-stack images was performed with Volocity software or with Fiji software and exported as 16-bit TIFF files.

### Assessment of stress fibre tilt

Angle (°) of stress fibre tilt was measured using images of actin cytoskeleton in cells labelled either by phalloidin or SpyFastAct. First, a maximum projection of the Z-stack images was performed with Fiji software and exported as 16-bit TIFF files. Using these projected images, orientation analysis was performed using the OrientationJ Vector Field plugin in Fiji. The window size was set to 5 pixels and the cubic spline method was used. The output orientation map in degree (°) was saved as a 32-bit TIFF file for further analysis using a custom MATLAB script. For analysis of each image, its orientation map and a corresponding binary mask were used. The mask was obtained by thresholding of the actin image to obtain micropattern or cell outline. The mask defined the region of interest, with pixels inside the mask considered for further analysis by the MATLAB script.

In the custom MATLAB script, a sliding-window approach was used to extract local actin vectors. Briefly, square windows of 20 pixels were sampled across the image with a stride of 18 pixels. Windows were retained if at least 80% of the area overlapped with the mask. Within each valid window, the local orientation angles were extracted and the mean of all local orientation angles found within the binary mask were calculated for each image to represent the value of stress fibre tilt per cell. The actin vector field was visualised by overlaying vectors on the corresponding fluorescence image. Each apolar vector was drawn centred at the window coordinates and colour-coded to distinguish positive (cyan; greater than 0°, rightward) and negative (yellow; less than or equal to 0°, leftward). This produced a spatial map of local orientations superimposed on the fluorescence image.

### Assessment of 2D-microtissue

Cells were seeded on rectangular or parallelogram micropatterns at a density of 1×10^5^ cells ml^-1^ for 20 min. The medium containing unattached cells was then replaced with fresh DMEM and 2D-microtissues were incubated for a total of 48 hours before cell fixation using 4% paraformaldehyde-PBS for 10 min. Phase-contrast and widefield images of 2D-microtissue were taken using either a 10× (0.3 NA, UPLFLN 10×, Olympus) or 20× air objective (0.45 NA, LUCPLFLN 20X, Olympus) on an Olympus IX81 inverted microscope, equipped with Andor Neo 5.5 sCMOS camera and light source (Lumencor SOLA SE Light Engine). Single plane images of phase contrast and DAPI channels were taken. For live 2D-microtissue imaging, cell nuclei were stained with 1 µg ml^-1^ Hoechst 33342 for 10 min. Phase-contrast and widefield images were taken every 10 mins for a total of 48 hours. Each image contained a single microtissue and these images were subsequently used for measurement of average nematic director angle.

### Computation of nematic director orientation in 2D-microtissue

The following procedures were performed using a custom MATLAB script as reported in our earlier work^23^. Briefly, identification and segmentation of individual 2D-microtissue using phase contrast images was done by performing a Wiener filter with a neighbourhood size of 20×20 pixels to remove image noise. This was followed by an entropy filter with a 3×3 pixels structural element and morphological opening with a 9×9 pixels structural element. Otsu binarisation was then performed to segment the image, the segmented area at the centre of the image was selected as the segmentation mask. The bounding box enclosing this segmented area serves to represent the dimensions of the microtissue.

Segmentation of the nuclei was achieved using NICK adaptive binarization. A Wiener filter using a neighbourhood size of 9×9 pixels was performed prior to segmentation. The concaved-point based splitting algorithm was used to separate any overlapping nuclei. Segmented objects above the size of 10000 pixels were removed as these corresponded to the background regions, while objects smaller than 500 pixels were also removed as these were either fragmented nuclei or noise regions that were segmented by chance. The number of nuclei in the bounding box was also counted.

Local cell orientation in the phase contrast image was calculated by obtaining the nematic director field as described in ref 59. Briefly the orientation tensor was obtained using OrientationJ implemented in Fiji and the nematic director was obtained using a window size of 60×60 µm^2^ and 70% overlap. The orientation of each director was defined relative to the long axis of the rectangular micropattern, or relative to the perpendicular to the base in the case of the parallelogram. Director orientations within the region of interest were then used to assess the overall alignment of the microtissue. For each microtissue, the mean nematic director angle was obtained by averaging the orientations of all directors within that culture.

### Determining orientation pattern of rectangular 2D-microtissue

Each 300×600 µm 2D-microtissue was characterised by an array of 8×16 nematic directors. To assess orientation pattern, each microtissue was divided into four quadrants of 75×150 µm, each comprising a 8×4 array of nematic directors. For each quadrant, the mean nematic director angle was calculated from all directors within the region, yielding four mean values per microtissue. Each quadrant was then assigned a symbol based on the value for the mean nematic director angle as follow: /for angles ≥3° to ≤87°, \ for angles ≥–87° to ≤–3°, | for angles >–3° to <3°, or – for angles ≥–90 to <–87° and >87° to ≤90°. See also Extended Data Fig. 4.

From these quadrant symbols, microtissue orientation patterns were classified as follows: (1) И represents chiral alignment along the bottom-left to top-right diagonal and is defined by either (i) all quadrants = “/”, (ii) at least three “/” with “–” in the top or bottom quadrant, or (iii) at least three “/” with a “|”. (2) N represents chiral alignment along the bottom-right to top-left diagonal and is defined by either (i) all quadrants = “\”, (ii) at least three “\” with “–” in the top or bottom quadrant, or (iii) at least three “\” with a “|”. (3) C represents opposing orientations between top (= “/”) and bottom (= “\”) halves or quadrants, together with or without “|” in the middle two quadrants. (4) Z represents top and bottom quadrants = “\”, together with “/” or “|” in the middle quadrants. (5) S represents top and bottom quadrants = “/”, together with “\” or “|” in the middle quadrants. (6)| represents all four quadrants = “|”, or three quadrants = “|” with either “/”, “\”, or “–” in the top or bottom quadrant.

### Statistics and Reproducibility

The numbers of samples (n) of individual cells and microtissues analysed for all the quantitative data are specified in the figure legends. No statistical method was used to predetermine sample size. All images are representative of at least three independent experiments. All supplementary videos show representative data from at least three independent experiments. Prism software (version 10.6.1; GraphPad Software, LLC.) was used for graph plotting. For frequency distribution, a bin width of 2.5° is used for all histograms, except for histograms representing stress fibre tilt which uses a bin width of 2°.

### Modelling

The details of two models of intercellular chiral interactions, each incorporating experimentally motivated assumptions about cell shape, stochastic motility, and alignments, are as follows. In both models, individual cells are approximated as rods, defined by their 2D centre-of-mass position *x* and orientation θ.

### Random movements

The cells undergo translational and rotational diffusion, accounting for intrinsic fluctuations in motility. The position ***x*** and orientation θ evolve via:

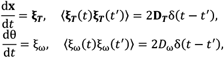

where ***ξ***_***T***_ and *ξ*_*ω*_ are white noise, ***D***_*T*_ is the translational diffusion tensor, and D_*ω*_ the rotational diffusion coefficient. Motivated by the high aspect ratio (ϵ) of the cells, we adopt a slender-body approximation, assuming enhanced diffusion along the long axis:

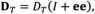

where *e* is the unit vector along the long axis, *I* is the identity matrix, *D*_*T*_ is the baseline diffusion in the direction perpendicular to *e*, and L is the cell length.

### Cell-cell and cell-boundary alignment

Both models incorporate nematic alignment interactions between cells, reflecting experimental observations that neighbouring cells tend to align their long axes. Inspired by the Kuramoto model^60^, we describe this alignment via a sinusoidal coupling angular velocity:

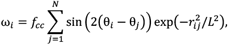

where N is the total number of cells, *r*_*ij*_ is the distance between cell i and j (nondimensionalized by *L*, the characteristic distance between the neighbouring cells, so the alignment is local), and *f*_*cc*_ is the alignment strength.

Interactions with the boundary are treated analogously: the boundary is modelled as a set of fixed “virtual cells” with orientation θ_*b*_ determined by the local boundary direction. The corresponding alignment angular velocity is:

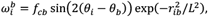

where *r*_*ib*_ is the distance between the cell and the nearest boundary segment, and *f*_*cb*_ is the cell-boundary alignment strength.

### Boundary-cell model: chirality originates at the boundary

In the boundary-cell model, we posit that chirality originates from asymmetric interactions between cells and the boundary. Motivated by experimental observations that cells near the edge tilt clockwise with respect to the boundary (Fig. 1c,d), we introduce a chiral tilt *α*_bc_ in the alignment interaction:

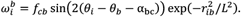

Here, *α*_bc_ denotes the intrinsic chiral tilt induced by stress fibre rotation. Based on experimental measurements, we set the mean value of *α*_bc_ to 30^°^, while its standard deviation (60^°^) was determined by fitting simulation outputs to experimental alignment statistics (Fig. 3). This directional bias enforces a persistent clockwise tilt at the boundary, which subsequently propagates into the tissue through cell–cell alignment.

### Inner-cell model: chirality originates in the bulk

The inner-cell model explores an alternative mechanism in which each cell develops intrinsic chirality independent of boundary effects. Experiments suggest that stress fibres in individual cells rotate clockwise, and that the cell long axis gradually reorients to align with them (Fig. 2). We denote ψ as the orientation of the stress fibre bundle, and model the coupled dynamics of θ and ψ via:

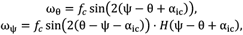

where *f*_*c*_ is the coupling strength (fit from experiments), and *H*(*x*) is the Heaviside step function enforcing clockwise-only rotation of ψ. This mechanism maintains a persistent offset angle *α*_ic_ between stress fibres and the cell axis, to reflect the experimental observation that when the cell orientation and shape are fixed on adhesive islands, the stress fibres tilt clockwise relative to the cell axis. While θ may fluctuate symmetrically, ψ rotates only clockwise, producing a net chiral drift over time. In this model, chirality emerges in the bulk of the micro-tissue and is subsequently stabilized by the boundary, inverting the direction of propagation relative to the boundary-cell model.

### Parameter summary

All parameters used in the simulations are listed in Table 1. Orders of magnitude for values for random movements and rotations, cell sizes and numbers were obtained directly from the experimental measurements. Alignment strengths were found by fitting simulation results to experimentally obtained orientation pattern statistics.

**Table 1:**
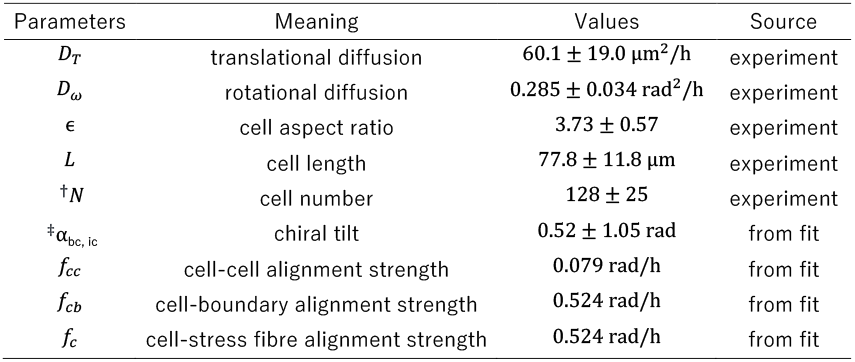
Model parameters and their sources. ^†^*N* scales with island area; reported value corresponds to a 600 × 300 μm^2^ rectangle. ^‡^*α*_bc, ic_ is sampled from a normal distribution.

## Notes

### Competing Interest Statement

The authors have declared no competing interest.

## References

1. Levin, M. Left-right asymmetry in embryonic development: a comprehensive review. Mech Dev 122, 3–25 (2005).

2. Henley, C.L. Possible Origins of Macroscopic Left-Right Asymmetry in Organisms. Journal of Statistical Physics volume 148, 741–775 (2012).

3. Naganathan, S.R., Middelkoop, T.C., Furthauer, S. & Grill, S.W. Actomyosin-driven left-right asymmetry: from molecular torques to chiral self organization. Curr Opin Cell Biol 38, 24–30 (2016).

4. Inaki, M., Sasamura, T. & Matsuno, K. Cell Chirality Drives Left-Right Asymmetric Morphogenesis. Front Cell Dev Biol 6, 34 (2018).

5. Hamada, H. & Tam, P. Diversity of left-right symmetry breaking strategy in animals. F1000Res 9 (2020).

6. Rahman, T., Zhang, H., Fan, J. & Wan, L.Q. Cell chirality in cardiovascular development and disease. APL Bioeng 4, 031503 (2020).

7. Lapraz, F., Fixary-Schuster, C. & Noselli, S. Brain bilateral asymmetry - insights from nematodes, zebrafish, and Drosophila. Trends Neurosci 47, 803–818 (2024).

8. Tsikolia, N., Nguyen, D.T.L. & Tee, Y.H. Mechanisms of left-right symmetry breaking across scales. Curr Opin Cell Biol 95, 102564 (2025).

9. Davison, A. Flipping Shells! Unwinding LR Asymmetry in Mirror-Image Molluscs. Trends Genet 36, 189–202 (2020).

10. Kuroda, R. & Abe, M. The pond snail Lymnaea stagnalis. Evodevo 11, 24 (2020).

11. Thurman, C.L., Alber, R.E., Hopkins, M.J. & Shih, H.T. Morphological and Genetic Variation Among Populations of the Fiddler Crab Minuca burgersi (Holthuis, 1967) (Crustacea: Brachyura: Ocypodidae) from Shores of the Caribbean Basin and Western South Atlantic Ocean. Zool Stud 60, e19 (2021).

12. Callaway, E. Rare ‘ambidextrous’ protein breaks rules of handedness. Nature 642, 278–279 (2025).

13. Hirokawa, N., Tanaka, Y., Okada, Y. & Takeda, S. Nodal flow and the generation of left-right asymmetry. Cell 125, 33–45 (2006).

14. Shinohara, K. & Hamada, H. Cilia in Left-Right Symmetry Breaking. Cold Spring Harb Perspect Biol 9 (2017).

15. Katoh, T.A. et al. Immotile cilia mechanically sense the direction of fluid flow for left-right determination. Science 379, 66–71 (2023).

16. Djenoune, L. et al. Cilia function as calcium-mediated mechanosensors that instruct left-right asymmetry. Science 379, 71–78 (2023).

17. Gros, J., Feistel, K., Viebahn, C., Blum, M. & Tabin, C.J. Cell movements at Hensen’s node establish left/right asymmetric gene expression in the chick. Science 324, 941–944 (2009).

18. Pfanzelter, J. et al. An active torque dipole across tissue layers drives avian left-right symmetry breaking. bioRxiv, 2025.2007.2016.665037 (2025).

19. Pieper, T.K. & Tsikolia, N. Left-right symmetry breaking: learning from the chicken. Frontiers in Cell and Developmental Biology Volume 13 - 2025 (2025).

20. Wan, L.Q. et al. Micropatterned mammalian cells exhibit phenotype-specific left-right asymmetry. Proc Natl Acad Sci U S A 108, 12295–12300 (2011).

21. Chen, T.H. et al. Left-right symmetry breaking in tissue morphogenesis via cytoskeletal mechanics. Circ Res 110, 551–559 (2012).

22. Tee, Y.H. et al. Cellular chirality arising from the self-organization of the actin cytoskeleton. Nat Cell Biol 17, 445–457 (2015).

23. Tee, Y.H. et al. Actin polymerisation and crosslinking drive left-right asymmetry in single cell and cell collectives. Nat Commun 14, 776 (2023).

24. Hozumi, S. et al. An unconventional myosin in Drosophila reverses the default handedness in visceral organs. Nature 440, 798–802 (2006).

25. Speder, P., Adam, G. & Noselli, S. Type ID unconventional myosin controls left-right asymmetry in Drosophila. Nature 440, 803–807 (2006).

26. Taniguchi, K. et al. Chirality in planar cell shape contributes to left-right asymmetric epithelial morphogenesis. Science 333, 339–341 (2011).

27. Lebreton, G. et al. Molecular to organismal chirality is induced by the conserved myosin 1D. Science 362, 949–952 (2018).

28. Davison, A. et al. Formin Is Associated with Left-Right Asymmetry in the Pond Snail and the Frog. Curr Biol 26, 654–660 (2016).

29. Kuroda, R. et al. Diaphanous gene mutation affects spiral cleavage and chirality in snails. Sci Rep 6, 34809 (2016).

30. Abe, M. & Kuroda, R. The development of CRISPR for a mollusc establishes the formin Lsdia1 as the long-sought gene for snail dextral/sinistral coiling. Development 146 (2019).

31. Middelkoop, T.C. et al. CYK-1/Formin activation in cortical RhoA signaling centers promotes organismal left-right symmetry breaking. Proc Natl Acad Sci U S A 118 (2021).

32. Noel, E.S. et al. A Nodal-independent and tissue-intrinsic mechanism controls heart-looping chirality. Nat Commun 4, 2754 (2013).

33. Ray, P. et al. Intrinsic cellular chirality regulates left-right symmetry breaking during cardiac looping. Proc Natl Acad Sci U S A 115, E11568–E11577 (2018).

34. Naganathan, S.R., Furthauer, S., Nishikawa, M., Julicher, F. & Grill, S.W. Active torque generation by the actomyosin cell cortex drives left-right symmetry breaking. Elife 3, e04165 (2014).

35. Jalal, S. et al. Actin cytoskeleton self-organization in single epithelial cells and fibroblasts under isotropic confinement. J Cell Sci 132 (2019).

36. Li, W. et al. Chiral growth of adherent filopodia. Biophys J 122, 3704–3721 (2023).

37. Kwong, H.K. et al. Cell chirality reversal through tilted balance between polymerization of radial fibers and clockwise-swirling of transverse arcs. eLife 12, RP92632 (2024).

38. Yamamoto, T. et al. Epithelial cell chirality emerges through the dynamic concentric pattern of actomyosin cytoskeleton. eLife 14, e102296 (2025).

39. Yamaguchi, A. et al. Class I myosins direct circumferential F-actin flows to define cell chirality. bioRxiv, 2025.2005.2006.648335 (2025).

40. Duclos, G. et al. Spontaneous shear flow in confined cellular nematics. Nat Phys 14, 728–732 (2018).

41. Maitra, A. & Lenz, M. Spontaneous rotation can stabilise ordered chiral active fluids. Nat Commun 10, 920 (2019).

42. Hoffmann, L.A., Schakenraad, K., Merks, R.M.H. & Giomi, L. Chiral stresses in nematic cell monolayers. Soft Matter 16, 764–774 (2020).

43. Li, X. & Chen, B. Dynamics of multicellular swirling on micropatterned substrates. Proc Natl Acad Sci U S A 121, e2400804121 (2024).

44. Rahman, T., Peters, F. & Wan, L.Q. Biomechanical Modeling of Cell Chirality and Symmetry Breaking of Biological Systems. Mechanobiol Med 2 (2024).

45. Parker, K.K. et al. Directional control of lamellipodia extension by constraining cell shape and orienting cell tractional forces. FASEB J 16, 1195–1204 (2002).

46. Chen, C.S., Alonso, J.L., Ostuni, E., Whitesides, G.M. & Ingber, D.E. Cell shape provides global control of focal adhesion assembly. Biochem Biophys Res Commun 307, 355–361 (2003).

47. Thery, M., Pepin, A., Dressaire, E., Chen, Y. & Bornens, M. Cell distribution of stress fibres in response to the geometry of the adhesive environment. Cell Motil Cytoskeleton 63, 341–355 (2006).

48. Isomursu, A. et al. Dynamic Micropatterning Reveals Substrate-Dependent Differences in the Geometric Control of Cell Polarization and Migration. Small Methods 8, e2300719 (2024).

49. Yashunsky, V. et al. Chiral Edge Current in Nematic Cell Monolayers. Physical Review X 12, 041017 (2022).

50. Guillamat, P., Blanch-Mercader, C., Pernollet, G., Kruse, K. & Roux, A. Integer topological defects organize stresses driving tissue morphogenesis. Nat Mater 21, 588–597 (2022).

51. Maroudas-Sacks, Y., Garion, L., Suganthan, S., Popović, M. & Keren, K. Confinement Modulates Axial Patterning in Regenerating Hydra. PRX Life 2, 043007 (2024).

52. Chen, S. et al. Chirality across scales in tissue dynamics. arXiv 2506.12276 (2025).

53. Chougule, A. et al. The Drosophila actin nucleator DAAM is essential for left-right asymmetry. PLoS Genet 16, e1008758 (2020).

54. Chin, A.S. et al. Epithelial Cell Chirality Revealed by Three-Dimensional Spontaneous Rotation. Proc Natl Acad Sci U S A 115, 12188–12193 (2018).

55. Zhang, H., Fan, J., Maclin, J.M.A. & Wan, L.Q. The Actin Crosslinker Fascin Regulates Cell Chirality. Adv Biol (Weinh) 7, e2200240 (2023).

56. Tamada, A., Kawase, S., Murakami, F. & Kamiguchi, H. Autonomous right-screw rotation of growth cone filopodia drives neurite turning. J Cell Biol 188, 429–441 (2010).

57. Stramer, B. & Mayor, R. Mechanisms and in vivo functions of contact inhibition of locomotion. Nat Rev Mol Cell Biol 18, 43–55 (2017).

58. Bruckner, D.B. et al. Learning the dynamics of cell-cell interactions in confined cell migration. Proc Natl Acad Sci U S A 118 (2021).

59. Tan, T.H. et al. Emergent chirality in active solid rotation of pancreas spheres. PRX Life 2, 033006 (2024).

## Supplementary References

60. Saw, T.B. et al. Topological defects in epithelia govern cell death and extrusion. Nature 544, 212–216 (2017).

61. Acebron, J.A. et al. The Kuramoto model: A simple paradigm for synchronization phenomena. Rev Mod Phys 77(1), 137 (2005).

